# Plasticity manifolds: Conjunctive changes in multiple ion channels mediate activity-dependent plasticity in hippocampal granule cells

**DOI:** 10.1101/747550

**Authors:** Poonam Mishra, Rishikesh Narayanan

## Abstract

The dentate gyrus (DG) was the first brain region to provide insights about synaptic and intrinsic plasticity. However, the assessment of intrinsic plasticity in DG has been surprisingly limited. We employed whole-cell patch-clamp recordings to explore the impact of behaviorally relevant theta-modulated burst firing, in the absence of synaptic stimulation, on intrinsic properties of rat DG granule cells. We found that theta burst firing induced a significant reduction in sub-threshold excitability and temporal summation, accompanied by an unexpectedly *contrasting* enhancement of action potential firing rate. We show that conjunctive changes in HCN, inward-rectifier potassium and persistent sodium channels mediated this form of plasticity, which was dependent on calcium influx through *L*-type calcium channels and inositol trisphosphate receptors. Our results unveil the expression of conjunctive plasticity in multiple channels, responding to the same activity pattern, establishing a *plasticity manifold* that could concomitantly mediate encoding and homeostasis in DG engram cells.

## INTRODUCTION

The concomitant roles of synaptic plasticity and neuron-specific intrinsic plasticity as putative cellular substrates of learning and memory are well established^1–6^. The DG has been implicated in spatial navigation, response decorrelation, pattern separation, engram formation, learning and memory^7–12^. The mechanisms behind and implications for synaptic plasticity to these physiological roles of the DG have been thoroughly investigated^7,10,12,13^. In contrast, the assessment of the protocols, mechanisms and implications associated with plasticity of intrinsic neuronal properties in the DG has been surprisingly limited^2,14^. This lacuna is especially striking because the first ever report of long-term potentiation (which was in the DG) also had reported a concomitant increase in granule cell excitability^12^, and because of prominent encoding roles for DG neurons as engram cells that could recruit neuron-specific changes to excitability^2,10^.

Here, seeking to fill this lacuna, we focused on identifying behaviorally relevant activity patterns that could induce intrinsic plasticity in DG granule cells and on mechanistically understanding such plasticity. In search of behaviorally relevant activity patterns, we first noted that theta frequency (4–10 Hz) oscillations are widely prevalent in the DG^15,16^ and granule cells exhibit theta-modulated bursts under *in vivo* conditions^17,18^. Although the impact of theta patterned stimuli has been widely studied with reference to synaptic plasticity^19^, the impact of theta burst firing in eliciting plasticity in neuronal intrinsic properties has not been explored. Therefore, employing whole-cell patch-clamp recordings, we explored the impact of intracellularly initiated theta-modulated burst firing, in the absence of synaptic stimulation, on intrinsic neuronal properties of DG granule cells.

We found that theta burst firing (TBF) induced a significant reduction in sub-threshold excitability and temporal summation, and an unexpected contrasting enhancement of supra-threshold excitability of DG granule cells. We provide strong lines of evidence, based on systematic analyses of signature electrophysiological characteristics and on results of experiments involving pharmacological agents, in support of conjunctive changes in HCN, inward-rectifier potassium and persistent sodium channels mediating this form of plasticity. Finally, we demonstrated that TBF-induced intrinsic plasticity was dependent on influx of cytosolic calcium, with *L-*type calcium channels and intracellular calcium stores contributing to calcium influx. Our results unveil the expression of conjunctive plasticity in multiple channels, responding to the *same* activity pattern, thereby establishing a *plasticity manifold* involving strong rules governing concomitant plasticity in different components. Such a plasticity manifold ensures that neural components have specific rules associated with how expression of plasticity in these components is co-regulated, thereby providing a molecular substrate for neurons to traverse across different stable states (say, each *encoding* different contexts without altering activity *homeostasis*) within the neuronal intrinsic manifold^4,20,21^. We postulate that the activity-dependent increase in neuronal firing rate could act as a putative substrate for the emergence of engram cells, and the associated reduction in sub-threshold excitability could concomitantly provide homeostatic balance to the encoding process.

## RESULTS

We performed whole-cell current clamp recordings from the somata of granule cells in the dentate gyrus. The behaviorally relevant activity pattern that we explored in our analyses is theta-burst firing (TBF), initiated by theta-patterned supra-threshold current injections (**Fig. 1c**, bottom) to induce action potential (AP) bursts (**Fig. 1c**, top). We first asked if intrinsic properties of DG granule cells change in response to behaviorally relevant activity patterns. We recorded several sub- and supra-threshold measurements before and after TBF induction to monitor changes in various intrinsic properties of granule cell (**Supplementary Table S1**).

**Figure 1:**
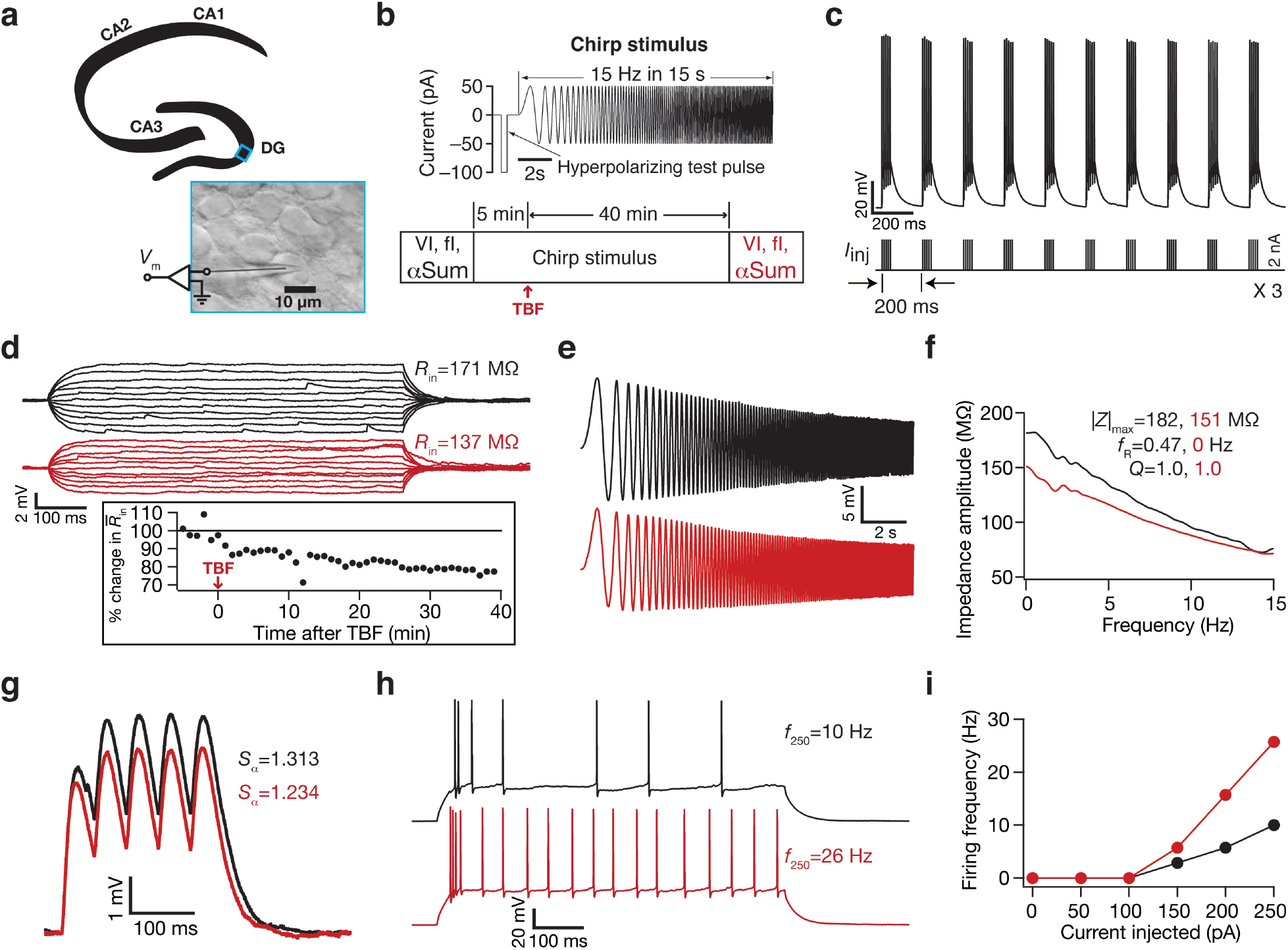
Illustration of theta burst firing inducing contrasting changes in sub- *vs.* supra-threshold excitability in a dentate gyrus granule cell. (**a**) Illustration showing whole-cell patch clamp recording from a dentate gyrus granule cell. (**b**) Experimental protocol, illustrating initial recordings of basic measurements after establishing a patch in whole-cell current-clamp configuration followed by five minutes recording of baseline measurements (*VI* and *fI* curves followed by alpha summation; shown in black) using chirp stimulus (*top*). After establishing 5 min of stable baseline, theta-patterned current pulses were injected into the neuron to elicit theta burst firing (TBF). Neuronal response properties were monitored for a further 40 minutes employing the chirp stimulus. At the end of 40 minutes, same sets of basic measurements (*VI* and *fI* curves followed by alpha summation; shown in red) were recorded to compare them with their baseline counterparts. A large 100-pA hyperpolarizing current pulse was provided before the chirp current to estimate input resistance 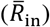 and to observe and correct series resistance changes through the course of the experiment. (**c**) Membrane voltage (*top*) recorded from the example granule cell in response to the theta-patterned current injection (*bottom*), showing 50 action potential elicited during a theta burst firing (TBF) pattern. This train was repeated thrice with 10 s inter-train intervals to form the TBF protocol employed in this study. (**d**) Voltage responses of an example neuron to 700 ms current pulses of amplitude varying from −25 pA to +25 pA (in steps of 5 pA), before (black) and 40 mins after (red) TBF. Input resistance (*R*_in_) was calculated as the slope of the plot depicting steady-state voltage response as a function of the injected current amplitude. Inset represents the temporal evolution of input resistance estimate 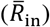 from neuronal response to the single hyperpolarizing pulse (depicted in panel **b**). (**e**) Voltage responses of the example neuron to the chirp current before (black: 0–5 min average) and 40 mins after (red: 40–45 min average) TBF. (**f**) Impedance amplitude computed from the current stimulus shown in panel **b** (*top*) and the voltage responses shown in panel **e**. |*Z*|_max_ represents the maximum impedance amplitude, *Q* is resonance strength and *f*_R_ denotes resonance frequency. (**g**) Voltage responses of the example neuron to 5 alpha-current injections arriving at 20 Hz, recorded before (black) and 40 mins after (red) TBF. Temporal summation ratio (*S*_α_) was computed as the ratio of the amplitude of the fifth response to that of the first. (**h**) Voltage response of the example neuron to a 700-ms current pulse of 250 pA, before (black) and 40 mins after (red) TBF. (**i**) Frequency of AP firing plotted as a function of injected current amplitude for the example cell. Note that these are firing frequencies converted from the number of spikes for the 700-ms injection duration. All traces and measurements shown in this example are from a single experiment.

### Theta burst firing elicited contrasting patterns of plasticity in sub- *vs.* supra-threshold physiological properties of granule cells

We found a significant long-term decrease in input resistance of GC post-TBF that persisted for more than 40 mins (**Fig. 1d, Fig. 2a–c**). The membrane potential also depolarized (~7 mV) through this period, but all measurements were obtained at the initial RMP to avoid confounds because of altered driving forces to different ion channels. Consistent with the reduction in *R*_in_, we also found a concomitant persistent reduction in impedance amplitude (**Fig. 1e–f**), which was quantified using maximal impedance amplitude (|*Z*|_max_; **Fig. 2d–e**), and in temporal summation (**Fig. 1g, Fig. 2g**) measured from the neuron’s response to five alpha excitatory postsynaptic currents. Despite these reductions in sub-threshold excitability measurements, which should have typically resulted in reduced AP firing^22,23^, we found an unexpected and contrasting enhancement in AP firing rate after TBF (**Fig. 1h–i, Fig. 2f**). This was coupled with changes in the firing patterns of neuron, whereby there was an occurrence of doublets at the beginning of the voltage response, quantified as a significant reduction in the first ISI after TBF (**Fig. 2g**). This form of contrasting plasticity was consistently reproducible (*n*=32; **Supplementary Table S1**) with measurements significantly different from 45 min control recordings (*n*=10) where no plasticity induction protocol was employed (**Fig. 2**).

**Figure 2.**
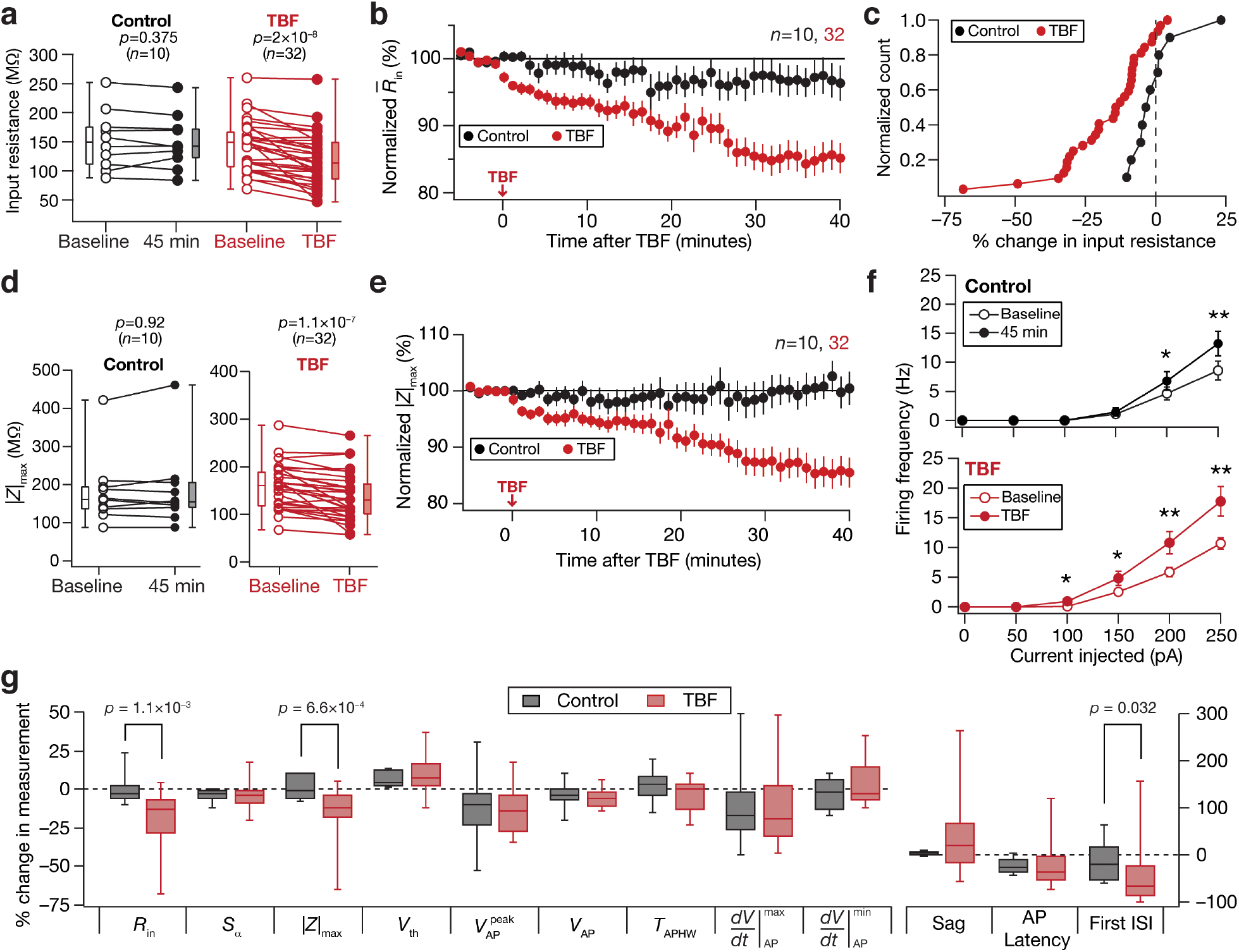
Theta burst firing induced contrasting changes in sub- *vs.* supra-threshold excitability in dentate gyrus granule cells. For all panels, the Control (black) group (*n*=10) corresponds to experiments where no protocol was applied through the 45-min period of the experiment, and TBF (red) group (*n*=32) is for neurons subjected to theta burst firing. For the TBF group, “Baseline” measurements were obtained before TBF and “TBF” measurements were after TBF. (**a**) Population data representing change in input resistance (*R*_in_) at the beginning (*empty circles*) and the end (*filled circles*) of the experiment. (**b**) Temporal evolution of percentage change in input resistance estimate (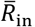; Mean ± SEM). (**c**) Normalized count of neurons from panel (**a**) plotted as functions of percentage change in *R*_in_ both for control and TBF case. (**d–e**) Same as (**a–b**), representing impedance amplitude (|*Z*|_max_). (**f**) Frequency of firing plotted as a function of injected current amplitude for both groups. *: *p*<0.05; **: *p*<0.005. Student’s *t* test. (**g**) Plots comparing percentage change in various measurements from their initial values to end of experiment values, for both control and TBF experiments. The list of symbols and corresponding measurements is enumerated in **Supplementary Table S1**. The Wilcoxon signed rank test was used for *p* value calculation in panels (**a**) and (**d**), for comparing measurements from the same set of cells. The Wilcoxon rank sum test was employed for *p* value calculation in panel (**g**), to compare percentage changes in the control group *vs*. those in the TBF group.

In one set of experiments, we performed physiological measurements before and after TBF at the respective RMP values *without* injecting current to maintain membrane potential at the initial value of the RMP (**Supplementary Fig. S1**). We found consistent TBF-induced reductions in input resistance and concomitant increases in firing rate, even when measurements were made at respective RMP values (**Supplementary Fig. S1**). Importantly, there was significant cell-to-cell variability in the amount of plasticity expressed in each of the different measurements we employed, pointing to the expression of *plasticity heterogeneities* in the granule cell population. However, heterogeneities in TBF-induced sub- or supra-threshold plasticity did not show strong correlations with heterogeneities in baseline neuronal intrinsic properties (**Supplementary Fig. S2**). Together, our results show that TBF yielded concomitant yet contrasting plasticity of sub- and supra-threshold intrinsic excitability in DG granule cells.

### TBF-induced plasticity in AP firing rate is consequent to a competition between changes in sub-threshold excitability and AP threshold

We quantified changes in 13 different sub- and supra-threshold measurements to assess the multifarious TBF-induced changes in granule cell physiology. Pairwise scatter plots of TBF-induced concomitant changes in these measurements revealed pronounced cell-to-cell variability in the expression of plasticity in each of these measurements (**Fig. 3**). Are heterogeneities in TBF-induced plasticity in these measurements related to each other, whereby specific pairs of measurements exhibited correlated changes despite each measurement manifesting heterogeneities? To assess this, we computed pairwise Pearson’s correlation between the TBF-induced percentage changes in each of the 13 individual measurements (**Fig. 3**). Among measurements that exhibited strong positive correlations in their TBF-induced changes were between measures of sub-threshold excitability (*R*_in_, |*Z*|_max_, *S*_α_) and between AP measurements (*V*_AP_, 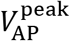, *V*_AP_, 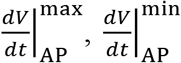). On the other hand, TBF-induced changes in AP half width showed strong negative correlations with each of changes in *V*_AP_, 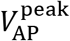, *V*_AP_, 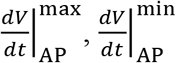, and changes in latency to first spike exhibited strong negative correlations with changes in AP threshold, and AP firing. Importantly, we noted strong correlations between changes in AP firing rate and changes in AP threshold, suggesting that changes in AP threshold could be mediating the enhanced firing rate despite a reduction in subthreshold excitability. The two groups of strongly correlated changes (Group 1: subthreshold measurements; Group 2: AP measurements), in conjunction with this strong relationship between AP firing and AP threshold changes suggested that TBF-induced changes in firing rate were regulated by two concurrent yet competing mechanisms. The first is a reduction in subthreshold excitability that reduces firing rate and the second is a hyperpolarizing shift in the AP threshold that would enhance AP firing. We noted that the absence of strong correlations (positive or negative) between changes in sub- *vs*. suprathreshold measurements could be consequent to this competition. Together, the contrasting changes introduced in sub- *vs*. supra-threshold measurements, in conjunction with the competition between subthreshold excitability and AP threshold in regulating firing rate pointed to conjunctive changes in multiple ion channels that mediated TBF-induced intrinsic plasticity in DG granule cells.

**Figure 3.**
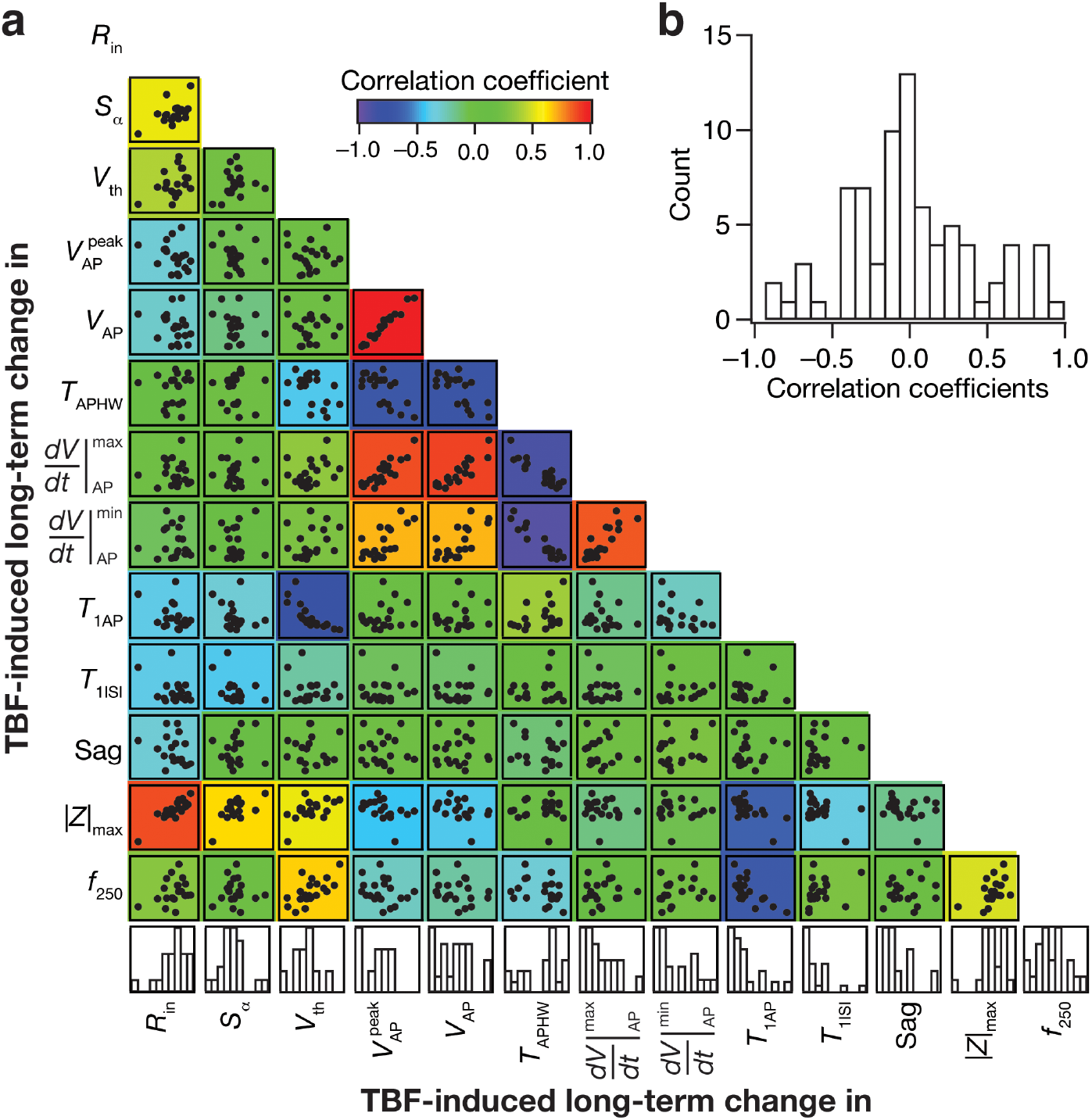
Differential pair-wise correlations among TBF-induced changes in thirteen electrophysiological measurements. (**a**) Pair-wise scatter plot matrices of TBF-induced changes in 13 sub- and supra-threshold measurements recorded from granule cells (*n*=20). These scatter plot matrices are overlaid on the corresponding color-coded correlation matrices. The bottom most row presents the histogram of percentage changes for corresponding measurements spanning their respective ranges. The list of symbols and corresponding measurements is enumerated in **Supplementary Table S1**. (**b**) Distribution of correlation coefficient values for 13 measurements corresponding to the pair-wise scatter plots shown in panel **a**.

### Signature changes in several physiological measurements point to changes in resting conductances as a candidate mechanism behind TBF-induced reduction in sub-threshold excitability

Focusing on TBF-induced changes in sub-threshold measurements, in our study, we observed a depolarizing shift in the resting membrane potential and reductions in input resistance, maximal impedance amplitude and temporal summation. These changes are broadly consistent with an increased current passage through HCN channels, which are hyperpolarization-activated channels that mediate an inward current that is active under resting conditions. Specifically, it has been shown that increases in HCN channels and the associated *h* current result in depolarization of resting membrane potential, a reduction of input resistance, impedance amplitude and temporal summation^23–25^.

Motivated by these observations, we asked if other TBF-induced changes in sub-threshold physiological measurements of granule cells were also consistent with changes in HCN channels. First, as HCN channels are slow conductances, their impact on neuronal excitability for relatively high-frequency inputs is low as the slow activation and deactivation profiles of these channels are inconsistent with such fast inputs. This has been employed^26^ as an effective mechanism to distinguish between changes in leak conductance (which would change excitability equally at all frequencies) or membrane capacitance (which would alter excitability preferentially at higher frequencies) or HCN channels (which would alter excitability preferentially at lower frequencies). To assess this, we quantified changes in impedance amplitude as a function of frequency, and asked if changes were predominantly at lower frequencies (**Fig. 4a–b**). We observed significantly higher TBF-induced changes in impedance amplitude at lower frequencies compared to changes in impedance amplitude at higher frequencies (**Fig. 4a–b**), again pointing to changes in HCN channels as a substrate for TBF-induced subthreshold changes. In contrast, changes in impedance amplitude were negligible in control experiments where no activity-dependent protocol was presented (**Fig. 4a–b**). These analyses point to the ability of impedance profiles to discern important underlying differences, owing to the frequency-dependent nature of the measurements. As neurons seldom receive steady-state inputs under ethological conditions, impedance constitutes a better measure of excitability than measures based on pulse-current injections^26^.

**Figure 4.**
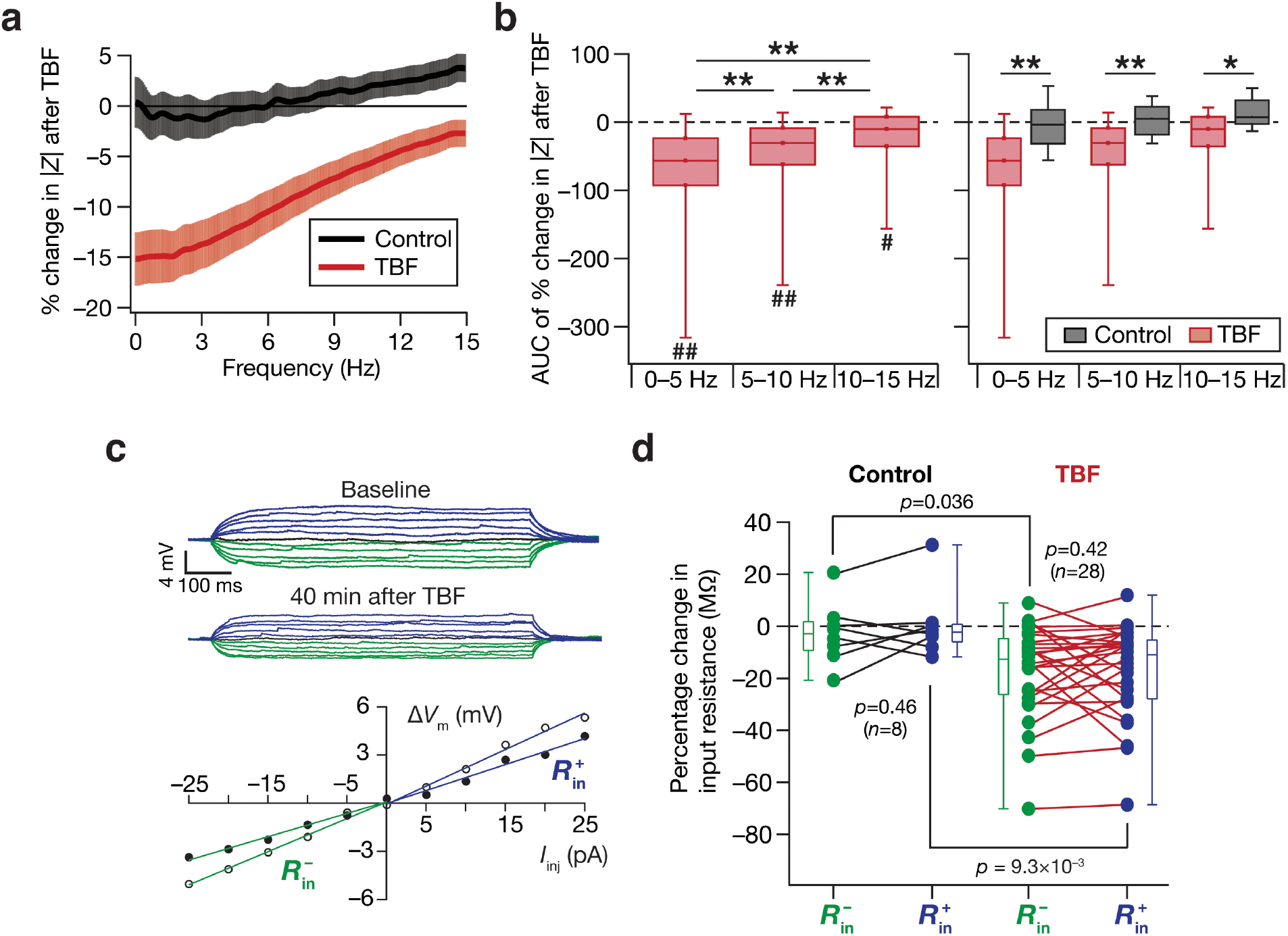
Analyses of changes in steady-state and frequency-dependent measurements point to HCN channels as a candidate mechanism behind TBF-induced reduction in sub-threshold excitability. (**a**) Percentage changes in impedance amplitude as a function of frequency for Control (no protocol) and TBF groups. In the control group, the percentage changes were computed as the change in impedance amplitude measured at 45-min, compared to the initial measurements. In the TBF group, the percentage changes are between those measured before and 40 mins after TBF. (**b**) Quantification of percentage changes in |*Z*| within three frequency bands (0–5 Hz, 5–10 Hz and 10–15 Hz). The value for each neuron were computed as the area under the curve (AUC) of percentage change in |*Z*|*. Left,* Plot showing the quartiles of AUC of percentage change in |*Z*| for neurons in the TBF group. The symbol “#” refers to the outcomes of Wilcoxon rank sum test on whether TBF-induced changes in |*Z*| within that frequency band were significantly different from zero. #: *p*<0.05; ##: *p*<0.001. The symbol “*” refers to the outcomes of Wilcoxon signed rank test, assessing whether TBF-induced changes in |*Z*| across the different frequency bands were significantly different from one another. **: *p*<0.001. *Right,* Plot showing the quartiles of AUC of percentage change in |*Z*| for neurons in the Control and TBF groups. The symbol “*” refers to the outcomes of Wilcoxon rank sum test, assessing whether TBF-induced changes in |*Z*| within each frequency band were significantly different between the Control and the TBF groups. *: *p*<0.05; **: *p*<0.005. Note that for the Control group, changes in |*Z*| within none of the three frequency bands were significantly different from zero, or significantly different from one another. Note that the representation has been split into two separate graphs to avoid clutter of symbols denoting statistical test outcomes. (**c**) Voltage responses of an example neuron to 700 ms current pulses of amplitude varying from −25 pA to +25 pA (in steps of 5 pA), recorded before (*Baseline*) and 40 mins after TBF. The responses colored blue and green are for positive and negative current injections, respectively. Input resistance values, computed from depolarizing 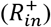 and hyperpolarizing 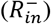 responses, were slopes of the plots depicting steady-state voltage responses as functions of positive and negative current injections, respectively (*bottom*). 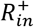 and 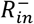 were calculated for traces obtained before (open circles) and after (closed circles) TBF (*bottom*). (**d**) Population level analysis to quantify percentage changes (at the end of experiment, compared to the measurement at the beginning) in 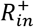 (blue) and 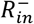 (green) independently for Control (*n*=8) and TBF (*n*=28) experiments. The *p* values presented for comparisons within each group (either Control or TBF) correspond to Wilcoxon signed rank test and *p* values across groups are for Wilcoxon rank sum test.

Our results thus far point to changes in a sub-threshold conductance that is active at rest, as all our measurements were obtained at the initial RMP. If this conductance that is active at rest were a depolarization-activated conductance, TBF-induced changes in voltage responses to depolarizing current pulses (where the channel is activated/deactivated) would be larger compared to responses to hyperpolarizing pulses (which is relatively beyond the channel’s activation/deactivation range). On the other hand, a hyperpolarization-activated conductance would show an opposite response profile, whereby there would be larger changes to hyperpolarized current injections compared to depolarized current injections. If the amount of changes were invariant to hyperpolarizing and depolarizing pulses, it would point to changes in a voltage-independent conductance or in a conductance whose activation/deactivation profile is linear within the range of measurement. To assess this, we calculated TBF-induced changes in two measures of input resistance, computed independently from voltage responses to depolarizing 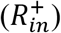 or hyperpolarizing 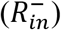 current injections (**Fig. 4c**). We performed this analysis both for control and TBF experiments and observed no significant difference between percentage changes in 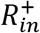 *vs.* 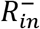 for either control or TBF experiments (**Fig. 4d**). However, consistent with prior observations on *R*_in_ (**Fig. 2**), the percentage changes in 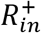 or 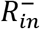 between control and TBF experiments were highly significant (**Fig. 4d**). These observations point to TBF-induced changes in a resting conductance whose activation profile is broadly linear in the range explored here, thereby ruling out the involvement of most depolarization-activated conductances.

Together, signature TBF-induced changes in several physiological measurements (**Supplementary Table S1**), — including depolarization in membrane potential, reductions in temporal summation (**Figs. 1–2**), dominant changes in excitability for inputs with lower frequencies (**Fig. 4a–b**) and consistent changes in input resistance for positive *vs.* negative pulse currents (**Fig. 4c–d**), — strongly point to a role for changes in HCN channels and other resting conductances as mechanisms underlying the observed reduction in sub-threshold excitability.

### Synergistic interactions between HCN and inward-rectifier potassium channels mediate TBF-induced reduction in sub-threshold excitability of DG granule cells

Encouraged by these strong lines of evidence on the potential role of resting conductances in sub-threshold plasticity, we first assessed the impact of two prominent resting conductances, HCN and inward-rectifier potassium (K_ir_) channels, on sub- and supra-threshold measurements in DG granule cells. We found that treatment with 20 μM ZD7288, a selective antagonist of HCN channels, resulted in significant *hyperpolarization* of RMP and significant increases in input resistance, impedance amplitude, temporal summation and action potential firing frequency (**Supplementary Fig. S3**). Treatment with 50 μM BaCl_2_, a selective blocker of K_ir_ channels, resulted in significant *depolarization* of RMP and significant increases in input resistance, impedance amplitude, temporal summation and action potential firing frequency (**Supplementary Fig. S4**). We confirmed the independent action of these two pharmacological agents by first treating DG granule cells with BaCl_2_, and subsequent treatment with the mixture of BaCl_2_ and ZD7288. These experiments confirmed our observations with these blockers individually (**Supplementary Fig. S3–S4**) and together showed that blockade of either HCN or barium-sensitive K_ir_ channels independently yield significant increases in sub- and supra-threshold excitability of DG granule cells (**Supplementary Fig. S5–S6**). However, treatment with 0.3 μM tertiapin-Q, a selective blocker of specific subtypes of inward-rectifier potassium channels, did not show significant changes in sub- or supra-threshold measurements of DG granule cells (**Supplementary Fig. S7**).

Having identified the presence of two resting conductances (HCN and K_ir_) with significant impact on sub-threshold excitability, we first tested the time-dependent effect of ZD7288 on intrinsic properties of DG granule cells. In these control experiments, we found a slow yet significant increase in input resistance over a period of 45 minutes (**Fig. 5a–c**), which also reflected as increases in maximal impedance amplitude (**Fig. 5d–e**). Next, we performed TBF experiments in the presence of ZD7288, and found *R*_in_ and |*Z*|_max_ to increase with time, with the strength of these increases across TBF experiments statistically insignificant when compared to control experiments performed in the presence of ZD7288 (**Fig. 5a–e, Fig. 5g**). These results demonstrated that TBF-induced reduction in sub-threshold excitability (**Fig. 2**) did not express in the presence of ZD7288. Importantly, TBF-induced enhancement in supra-threshold excitability, manifesting as increased firing rate was not blocked by the presence of ZD7288 (**Fig. 5f**). To further confirm our conclusions on the role of HCN channels in TBF-induced plasticity, we performed long-term controls and TBF experiments in the presence of 20 μM zatebradine, a structurally distinct HCN-channel blocker. Consistent with our results with ZD7288, we found that TBF-induced reduction in sub-threshold excitability did not express, but TBF-induced enhancement in supra-threshold excitability was not affected, in the presence of zatebradine (**Supplementary Fig. S8**).

**Figure 5.**
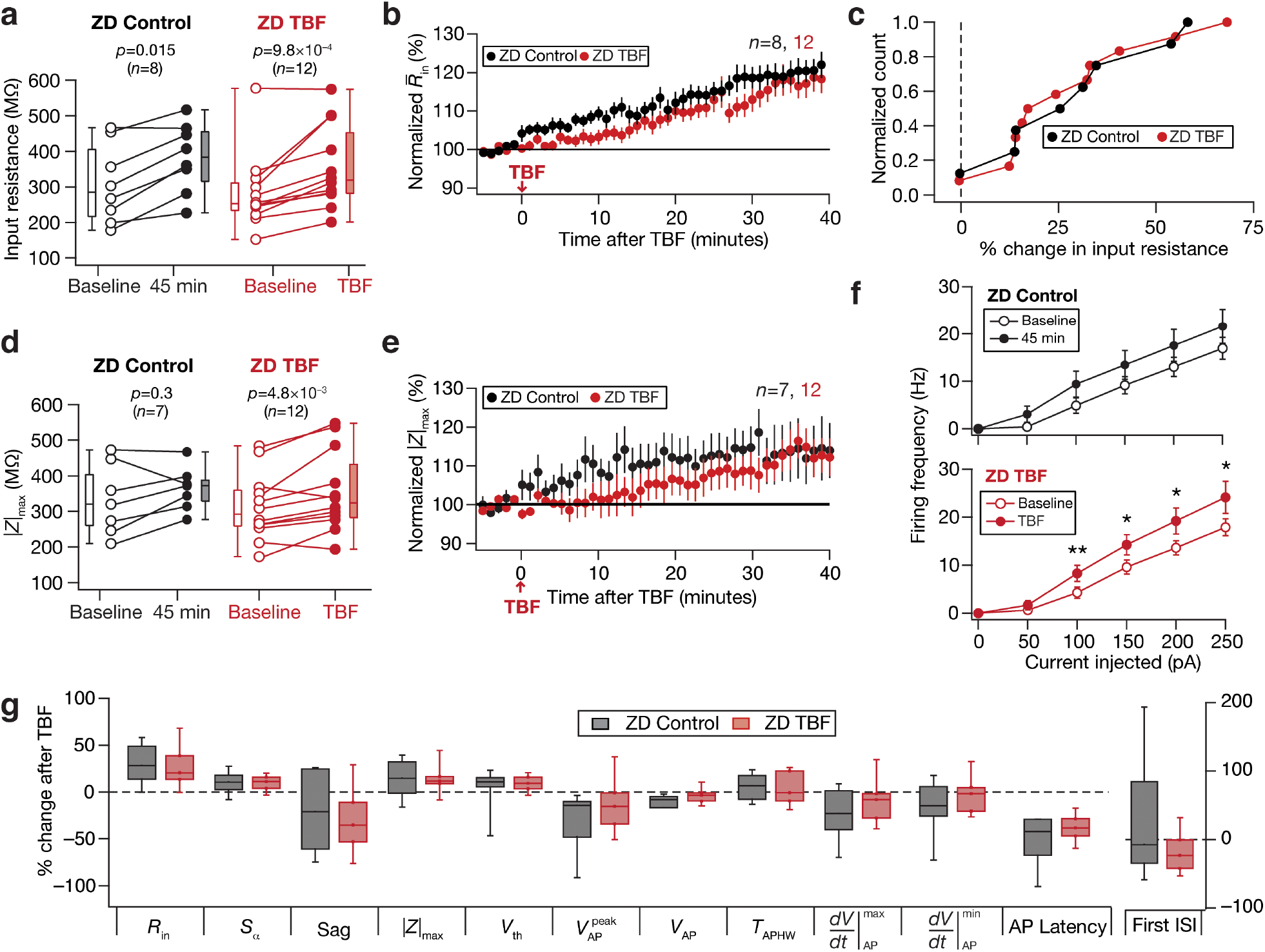
Activity-dependent reduction in sub-threshold excitability was blocked in the presence of ZD7288, a HCN channel blocker. For all panels, the ZD Control (black) group (*n*=8) corresponds to experiments where no protocol was applied through the 45-min period of the experiment, and ZD TBF (red) group (*n*=12) is for neurons subjected to theta burst firing. For the ZD TBF group, “Baseline” measurements were obtained before TBF and “TBF” measurements were after TBF. All experiments reported in this figure were performed in the presence of 20 μM ZD7288 in the bath and in the pipette solution. (**a**) Population data representing change in input resistance (*R*_in_) at the beginning (*empty circles*) and the end (*filled circles*) of the experiment. (**b**) Temporal evolution of percentage change in input resistance estimate (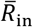; Mean ± SEM). (**c**) Normalized count of neurons from panel **a** plotted as functions of percentage change in *R*_in_. (**d–e**) Same as **a–b**, representing impedance amplitude (|*Z*|_max_). (**f**) Frequency of firing plotted as a function of injected current amplitude for both groups. *: *p*<0.05; **: *p*<0.005. Student’s *t* test. (**g**) Plots comparing percentage change in various measurements from their initial values to end of experiment values. The list of symbols and corresponding measurements is enumerated in **Supplementary Table S1**. The Wilcoxon signed rank test was used for *p* value calculation in panels **a** and **d**, for comparing measurements from the same set of cells. The Wilcoxon rank sum test was employed for *p* value calculation in panel **g**, to compare percentage changes in the control group *vs*. those in the TBF group.

Turning to K_ir_ channels, we performed long-term control experiments and TBF experiments in presence of 50 μM BaCl_2_. We found that the presence of BaCl_2_ blocked TBF-induced reduction in input resistance and other sub-threshold measurements (**Fig. 6**). Although the overall firing rate after TBF was higher than the respective baseline firing rates (**Fig. 6f;** BaCl_2_ TBF), we found the percentage change in AP threshold observed during TBF experiments to be significantly lower than the percentage change in AP threshold observed during control experiments (**Fig. 6g**). These experiments demonstrated that barium-sensitive channels play a critical role in mediating TBF-induced reduction in sub-threshold measurements.

**Figure 6.**
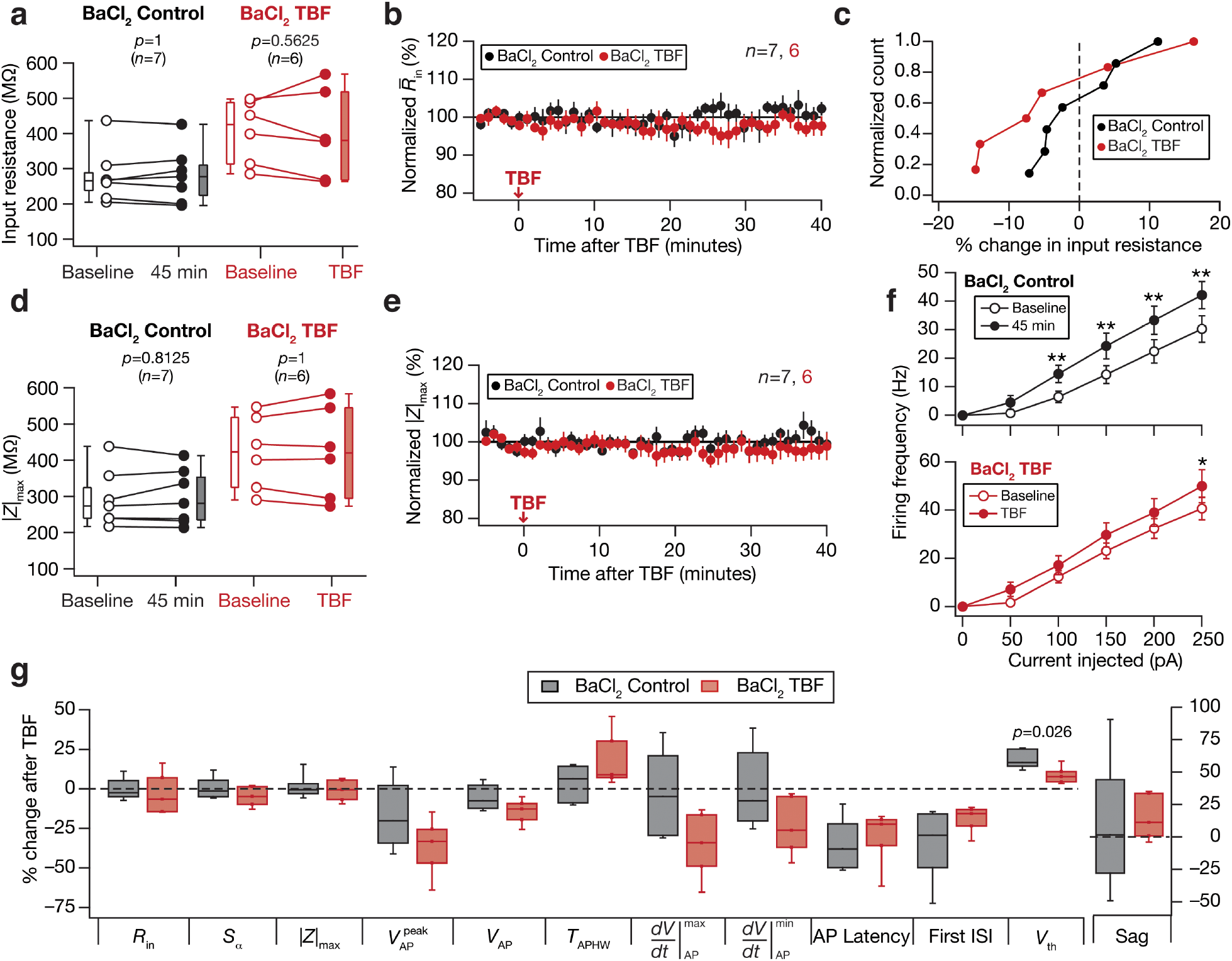
Activity-dependent reduction in sub-threshold excitability was blocked in the presence of barium chloride (BaCl_2_), a blocker of inward rectifier potassium channels. For all panels, the BaCl_2_ Control (black) group (*n*=7) corresponds to experiments where no protocol was applied through the 45-min period of the experiment, and BaCl_2_ TBF (red) group (*n*=6) is for neurons subjected to theta burst firing. For the BaCl_2_ TBF group, “Baseline” measurements were obtained before TBF and “TBF” measurements were after TBF. All experiments reported in this figure were performed in the presence of 50 μM BaCl_2_ in the bath. (**a**) Population data representing change in input resistance (*R*_in_) at the beginning (*empty circles*) and the end (*filled circles*) of the experiment. (**b**) Temporal evolution of percentage change in input resistance estimate (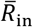; Mean ± SEM). (**c**) Normalized count of neurons from panel **a** plotted as functions of percentage change in *R*_in_. (**d–e**) Same as **a–b**, representing impedance amplitude (|*Z*|_max_). (**f**) Frequency of firing plotted as a function of injected current amplitude for both groups. *: *p*<0.05; **: *p*<0.005. Student’s *t* test. (**g**) Plots comparing percentage change in various measurements from their initial values to end of experiment values. The list of symbols and corresponding measurements is enumerated in **Supplementary Table S1**. The Wilcoxon signed rank test was used for *p* value calculation in panels **a** and **d**, for comparing measurements from the same set of cells. The Wilcoxon rank sum test was employed for *p* value calculation in panel **g**, to compare percentage changes in the control group *vs*. those in the TBF group.

Together, our physiological analyses (**Figs. 3–4**) and TBF experiments in the presence of blockers of HCN or K_ir_ channels (**Figs. 5–6; Supplementary Figs. S8**) provided lines of evidence for a critical role for synergistic interactions between these two resting conductances in mediating TBF-induced changes in subthreshold excitability. Whereas TBF-induced changes in subthreshold excitability could be explained either by increases in HCN or K_ir_ channels, TBF-induced *depolarization* in membrane potentials can be explained by an increase in the inward HCN, but not the outward K_ir_ channels (*cf.* **Supplementary Figs. S3–S5**). Therefore, it is possible that while changes in both HCN or K_ir_ channels synergistically contribute to the reduction in sub-threshold excitability, a relative dominance of changes to HCN channels results in TBF-induced depolarization of resting membrane potential. In addition, consistent with our previous analyses based on measurement correlations (**Fig. 3**), the dissociation in blockade of sub- *vs.* supra-threshold measurements by ZD7288 (**Fig. 5**) or zatebradine (**Supplementary Fig. S8**) pointed to the role of another channel conductance in mediating TBF-induced supra-threshold excitability.

### Plasticity in NaP channel results in TBF-induced increase in supra-threshold excitability of DG granule cells

Our observations on TBF-induced enhancement in supra-threshold excitability pointed to changes in a regenerative conductance. Motivated by the expression of a persistent sodium (NaP) channel in DG granule cells predominantly in axon initial segments^27–30^, and the role of these channels in regulating AP firing properties in several neuronal subtypes^31^, we first assessed a role for NaP channel in regulating DG granule cell excitability. We performed whole-cell current clamp recordings, measured baseline sub- and supra-threshold properties of DG granule cells, and then repeated these measurements in the presence of 20 μM riluzole (**Supplementary Fig. S9–S10**), an established inhibitor of persistent sodium currents^32^. We found that riluzole introduced no significant changes to resting membrane potential, sub-threshold excitability across different voltages or temporal summation (**Supplementary Fig. S9–S10**). In contrast, there was a pronounced reduction in firing rate of granule cells after application of riluzole (**Supplementary Fig. S9k–l**), without significant changes to AP properties (**Supplementary Fig. S10**). These results provided evidence for a role of persistent sodium channels in the regulation of supra-threshold excitability in DG granule cells.

We performed the long-term (45 min) control experiments in presence of riluzole without the theta burst firing protocol to examine any time dependent changes introduced by riluzole in intrinsic measurements. Although riluzole didn’t alter input resistance with acute treatment (**Supplementary Fig. S9**), we observed a significant reduction in input resistance over a 45-min period (**Fig. 7a–c**). This was observed even when we pre-treated the slice with riluzole before beginning our recordings. However, consistent with our observation under acute treatment of riluzole (**Supplementary Fig. S9**), impedance amplitude didn’t change over the course of the period in control experiments (**Fig. 7d–f**). We now performed TBF experiments in the presence of riluzole and compared these outcomes with riluzole control experiments (**Fig. 7**). We confirmed that the TBF protocol elicited 150 APs through the protocol period (**Supplementary Fig. S10n**), and there was no difference in AP firing pattern in the presence of riluzole. We found that TBF-induced reduction in input resistance was abolished in the presence of riluzole (**Fig. 7b–c**). In addition, analysis of |*Z*|_max_ in riluzole experiments without and with TBF presented clear evidence for the absence of any change in subthreshold excitability after TBF in the presence of riluzole (**Fig. 7d–f**). More importantly, the firing rate changes induced by TBF (**Fig. 2f**) was completely abolished in presence of riluzole with no further increase in firing rate post-TBF (**Fig. 7g**). We found that riluzole abolished TBF-induced changes in all sub-threshold measurements (**Fig. 7h**), as no significant difference was observed between control and TBF groups in the presence of riluzole. In this case, we could not quantify changes in AP properties as neurons did not fire at 250 pA for most of the recordings (both control and TBF groups).

**Figure 7.**
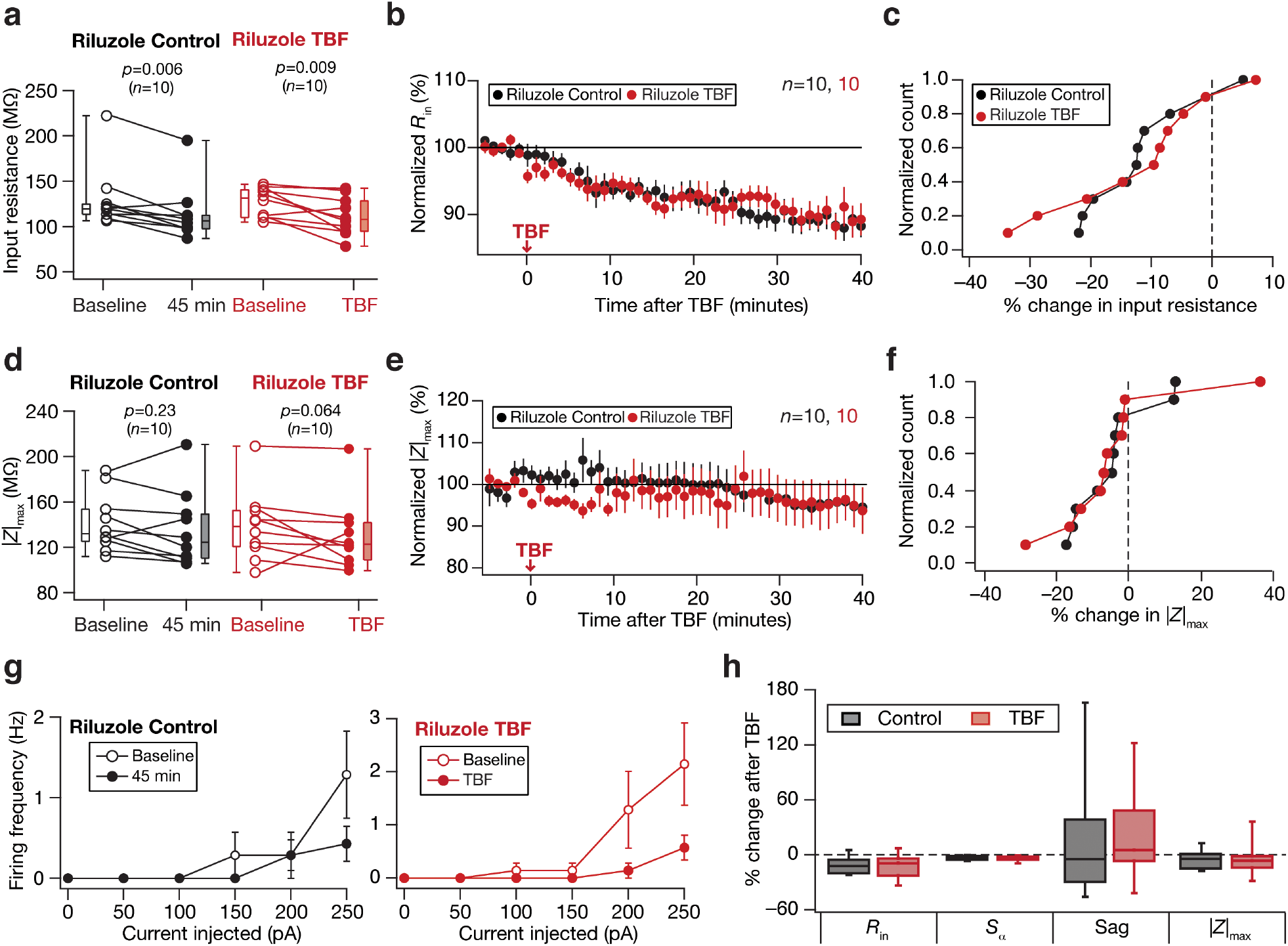
Activity-dependent intrinsic plasticity was blocked by Riluzole, a persistent sodium channel blocker. For all panels, the Riluzole Control (black) group (*n*=10) corresponds to experiments where no protocol was applied through the 45-min period of the experiment, and Riluzole TBF (red) group (*n*=10) is for neurons subjected to theta burst firing. For the Riluzole TBF group, “Baseline” measurements were obtained before TBF and “TBF” measurements were after TBF. All experiments reported in this figure were performed in the presence of 20 μM Riluzole in the bath. (**a**) Population data representing change in input resistance (*R*_in_) at the beginning (*empty circles*) and the end (*filled circles*) of the experiment. (**b**) Temporal evolution of percentage change in input resistance estimate (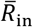; Mean ± SEM). (**c**) Normalized count of neurons from panel **a** plotted as a function of percentage change in *R*_in_. (**d–f**), Same as **a**–**c**, representing impedance amplitude (|*Z*|_max_). (**g**) Frequency of firing plotted as a function of injected current amplitude for both groups. *: *p*<0.05; **: *p*<0.005. Student’s *t* test. (**h**) Plots comparing percentage change in various sub-threshold measurements from their initial values to end of experiment values. The list of symbols and corresponding measurements are enumerated in **Supplementary Table S1**. The Wilcoxon signed rank test was used for *p* value calculation in panels **a** and **d**, for comparing measurements from the same set of cells. The Wilcoxon rank sum test was employed for *p* value calculation in panel **g**, to compare percentage changes in the control group *vs*. those in the TBF group.

Together, the direction of changes in several physiological measurements (**Figs. 2–4**), TBF experiments with ZD7288 (**Fig. 5**), BaCl_2_ (**Fig. 6**) or riluzole (**Fig. 7**) provided lines of evidence for conjunctive changes in HCN, inward-rectifier potassium and persistent sodium channels in mediating the contrasting TBF-induced plasticity profiles observed in sub- *vs*. supra-threshold changes in DG granule cells.

### Activity-dependent intrinsic plasticity in DG granule cells is calcium dependent

Several forms of neuronal plasticity are dependent on the prominent second messenger calcium. The magnitude and dynamics of calcium entry into the cytosol are well-established regulators of distinct signaling cascades, with cooperation and competition across different downstream cascades yielding different directions and strengths of plasticity in disparate neural components. Therefore, we asked if TBF-induced plasticity in DG granule cells was indeed dependent on calcium influx into the cytosol. To investigate this, we performed a set of control and TBF experiments in the presence of 30 mM BAPTA, a fast calcium chelator, in the patch pipette. We found no significant change in any measurement during control experiments in the presence of BAPTA in the pipette (**Fig. 8**). Importantly, when we monitored TBF-induced changes in the presence of BAPTA, we observed no significant activity-dependent change in any of the sub- or supra-threshold measurements (**Fig. 8**). We noticed that the overall firing rate of the cell was significantly high in presence of BAPTA in the pipette in both control as well as TBF experiments (**Fig. 8e** compared to **Fig. 2f**), pointing to a critical role of calcium-activated potassium channels in suppressing the excitability of DG granule cells^33^.

**Figure 8.**
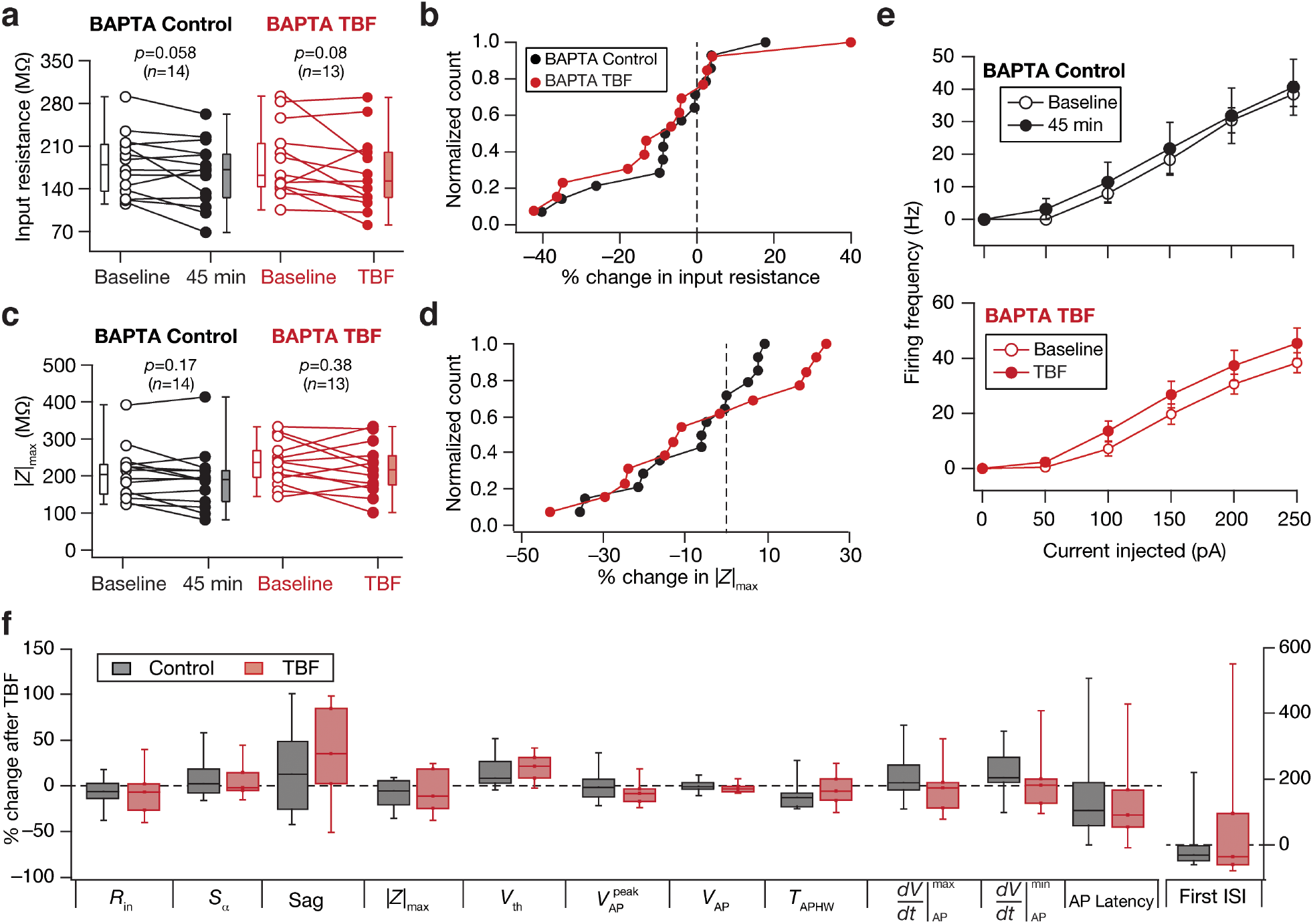
Activity-dependent intrinsic plasticity was blocked by BAPTA, a fast calcium chelator. For all panels, the BAPTA Control (black) group (*n*=14) corresponds to experiments where no protocol was applied through the 45-min period of the experiment, and BAPTA TBF (red) group (*n*=13) is for neurons subjected to theta burst firing. For the BAPTA TBF group, “Baseline” measurements were obtained before TBF and “TBF” measurements were after TBF. All experiments reported in this figure were performed in the presence of 30 mM BAPTA in the pipette solution. (**a**) Population data representing change in input resistance (*R*_in_) at the beginning (*empty circles*) and the end (*filled circles*) of the experiment. (**b**) Normalized count of neurons from panel **a** plotted as functions of percentage change in *R*_in_. (**c–d**) Same as **a–b**, representing impedance amplitude (|*Z*|_max_). (**e**) Frequency of firing plotted as a function of injected current amplitude for both groups. *: *p*<0.05; **: *p*<0.005. Student’s *t* test. (**f**) Plots comparing percentage change in various sub-threshold measurements from their initial values to end of experiment values. The list of symbols and corresponding measurements are enumerated in **Supplementary Table S1**. The Wilcoxon signed rank test was used for *p* value calculation in panels **a** and **c**, for comparing measurements from the same set of cells. The Wilcoxon rank sum test was employed for *p* value calculation in panel **g**, to compare percentage changes in the control group *vs*. those in the TBF group.

### Activity-dependent intrinsic plasticity is invariant to blockade of synaptic receptors

The plasticity induction protocol employed in this study for expression of intrinsic changes does not involve stimulation of synaptic receptors as the theta burst current pattern was injected directly into the soma. But, it is possible that different receptors driven by spontaneous activity could have interacted with intrinsic firing to play a role in plasticity induction^22^. To test this, we performed TBF experiments in the presence of AMPAR, GABA_A_R and GABA_B_R antagonists (**Fig. 9a–c, Supplementary Fig. S11**; Synaptic blockers). We found that the presence of synaptic blockers did not alter expression of TBF-induced plasticity (**Fig. 9a–c, Supplementary Fig. S11**; Synaptic blockers), although there was quantitatively less supra-threshold plasticity especially for lower current injections (**Fig. 9c**; Synaptic blockers. *cf.* **Fig. 2f**, TBF).

**Figure 9.**
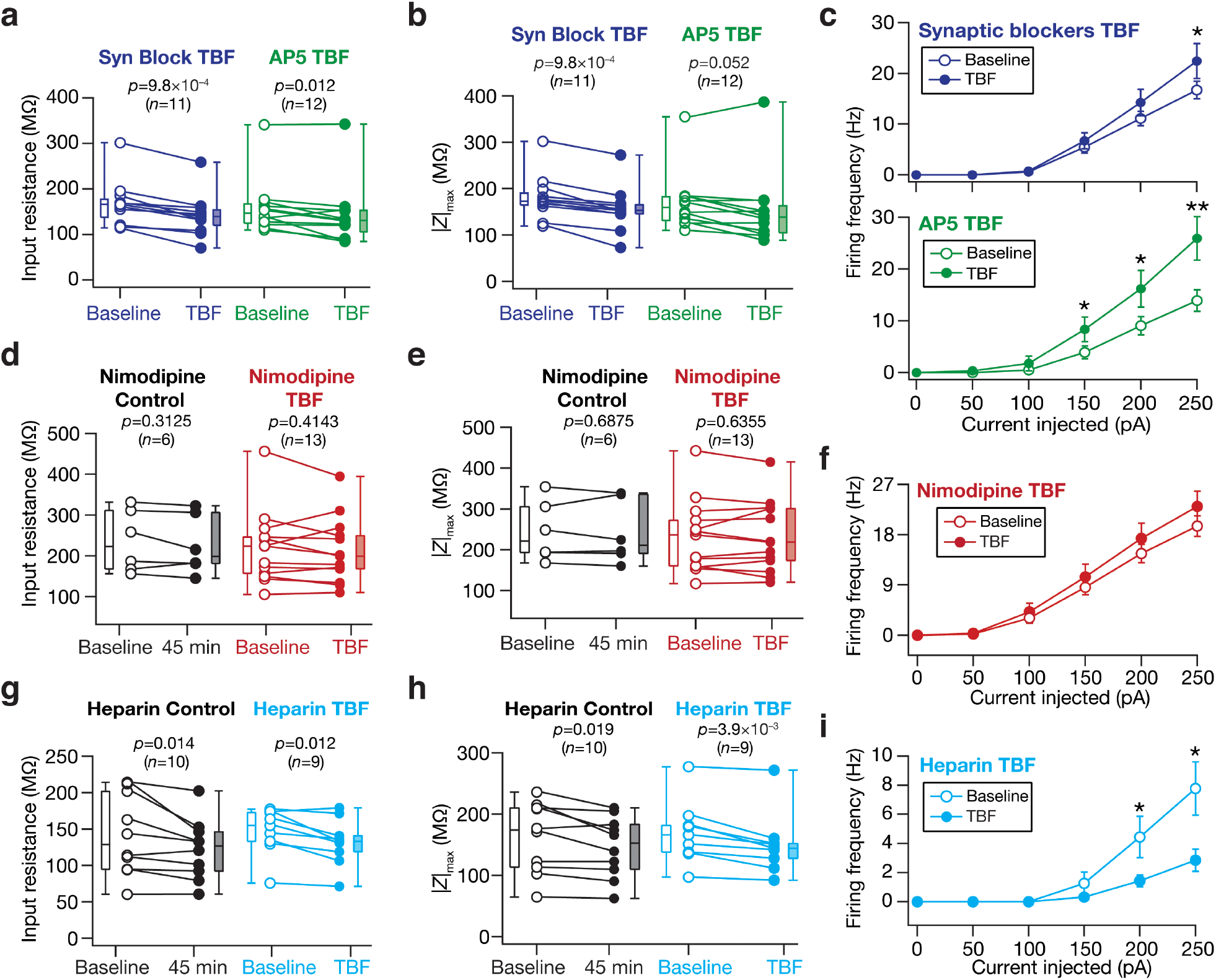
Activity-dependent intrinsic plasticity was independent of synaptic receptors, but was dependent on *L*-type calcium channels and InsP_3_ receptors. For all panels, “Baseline” measurements were obtained before TBF and “TBF” measurements were after TBF. (**a–c**) The “Syn Block” (blue) group (*n*=11) corresponds to experiments where neurons were subjected the TBF protocol in the presence of blockers of synaptic receptors (AMPAR, GABA_A_R and GABA_B_R) in the bath. The “AP5” (green) group (*n*=12) corresponds to experiments where neurons were subjected the TBF protocol in the presence of 50 μM AP5, an NMDAR antagonist, in the bath. Population data representing change in input resistance (*R*_in_; **a**), maximal impedance amplitude (|*Z*|_max_; **b**) and neuronal firing rate (**c**) at the beginning (*empty circles*) and the end (*filled circles*) of the experiment. See **Supplementary Fig. S11** for other plots associated with these experiments. (**d–f**) All experiments reported in these panels were performed in the presence of 10 μM Nimodipine in the bath. The Nimodipine Control (black) group (*n*=6) corresponds to experiments where no protocol was applied through the 45-min period of the experiment, and Nimodipine TBF (red) group (*n*=13) is for neurons subjected to theta burst firing. Population data representing change in input resistance (*R*_in_; **d**), maximal impedance amplitude (|*Z*|_max_; **e**) and neuronal firing rate (**f**) at the beginning (*empty circles*) and the end (*filled circles*) of the experiment. See **Supplementary Fig. S12** for other plots associated with these experiments. (**g–i**) All experiments reported in these panels were performed in the presence of 1 mg/mL Heparin in the pipette. The Heparin Control (black) group (*n*=10) corresponds to experiments where no protocol was applied through the 45-min period of the experiment, and Heparin TBF (cyan) group (*n*=9) is for neurons subjected to theta burst firing. Population data representing change in input resistance (*R*_in_; **g**), maximal impedance amplitude (|*Z*|_max_; **h**) and neuronal firing rate (**i**) at the beginning (*empty circles*) and the end (*filled circles*) of the experiment. See **Supplementary Fig. S13** for other plots associated with these experiments. The Wilcoxon signed rank test was used for *p* value calculation in panels **a**–**b**, **d**–**e** and **g**–**h**, for comparing measurements from the same set of cells. For firing rate plots, *: *p*<0.05; **: *p*<0.005. Student’s *t* test.

What component provided calcium influx into the cytosol that was essential for plasticity induction (**Fig. 8**)? We hypothesized NMDAR as a potential source of calcium, whereby spontaneous activity in conjunction with AP firing could have resulted in influx of calcium into the cytosol^22^. To test the hypothesis, we performed TBF experiments in presence of AP5, an NMDAR antagonist, and found TBF-induced sub- and supra-threshold plasticity to be independent of NMDARs (**Fig. 9a–c, Supplementary Fig. S11**; AP5), although TBF-induced changes in AP firing (**Fig. 9c**; AP5. *cf.* **Fig. 2f**, TBF) and certain AP properties (**Supplementary Fig. S11e**) were quantitatively higher in the presence of AP5.

### Activity dependent intrinsic plasticity is mediated by calcium influx through *L*-type calcium channels and inositol trisphosphate receptors

We hypothesized that repetitive firing triggered by the TBF protocol could recruit voltage-gated calcium channels (VGCCs) expressed in the granule cells as a source for calcium influx into the cytosol. To test this, we performed TBF experiments in presence of nimodipine, a blocker of *L*-type calcium channels, and compared the results with the control recordings (with nimodipine in the bath, but without TBF). We found that TBF-induced changes in both the sub- (**Fig. 9d–e, Supplementary Fig. S12a–c, e**) and supra-threshold (**Fig. 9f, Supplementary Fig. S12d–e**) measurements were abolished in presence of nimodipine. Specifically, in the presence of nimodipine, TBF-induced reduction in both input resistance (**Fig. 9d, Supplementary Fig. S12a–b**) and |*Z*|_max_ (**Fig. 9e, Supplementary Fig. S12c**) were abolished, apart from the absence of the associated increase in AP firing rate (**Fig. 9f**).

Finally, although the extracellular fluid is a prominent source for cytosolic calcium, calcium influx into the cytosol could also be through calcium channels expressed on the membrane of the endoplasmic reticulum (ER). Motivated by the results obtained from nimodipine experiments and expression of inositol trisphosphate (InsP_3_) receptors in DG granule cells^34–37^, we asked if ER calcium stores could also be a source of cytosolic calcium. We performed TBF in the presence of heparin, an established blocker of InsP_3_ receptors. We found that TBF-induced changes in various sub- and supra-threshold measurements were abolished in presence of heparin, providing evidence for ER calcium stores as a potential source for the induction of this form of plasticity (**Fig. 9g–i, Supplementary Fig. S13**).

Together, these experiments established that synergistic interactions between *L*-type calcium channels on the plasma membrane and InsP_3_ receptors on the ER membrane resulted in cytosolic calcium influx that mediated TBF-induced intrinsic plasticity.

## DISCUSSION

In this study, we showed that the behaviorally relevant theta-burst firing protocol reliably induced long-term intrinsic plasticity in dentate gyrus granule cells. We demonstrated that this intrinsic plasticity involved contrasting changes in sub- and supra-threshold measurements, with a reduction in sub-threshold excitability accompanying an enhancement in supra-threshold excitability. We reasoned that this form of activity-dependent plasticity should recruit changes in multiple ion channels, on the basis of our analyses involving a constellation of physiological measurements that changed with TBF. Through a combination of experiments involving pharmacological agents and analyses of physiological changes induced by TBF, we provide strong lines of evidence for this form of plasticity to involve conjunctive changes in multiple ion channels. Specifically, we presented evidence that synergistic interactions between plasticity in HCN and inward-rectifier potassium channels mediated TBF-induced reduction in sub-threshold excitability whereas plasticity in persistent sodium channels resulted in the accompanying increase in supra-threshold excitability. The calcium-dependence of TBF-induced intrinsic plasticity was evidenced from the blockade of plasticity by cytosolic BAPTA, a fast calcium chelator. Although TBF-induced intrinsic plasticity persisted in the presence of synaptic blockers (including NMDARs), nimodipine and cytosolic heparin independently blocked plasticity, suggesting the involvement of *L*-type calcium channels and inositol trisphosphate receptors as calcium sources.

### Activity-dependent intrinsic plasticity in DG granule cells and engram cell formation

Our study presents an important observation that DG granule cells are capable of expressing activity-dependent intrinsic plasticity *without* synaptic stimulation. We also demonstrate contrasting plasticity in sub- *vs.* supra-threshold excitability as a consequence of conjunctive changes in three distinct ion channel subtypes. The expression of cell-autonomous intrinsic plasticity expands the available repertoire of cellular substrates for information storage, from the perspective of what all can change in response to activity patterns^2–4,14^. These observations also extend to the formation of “engram cells”, whereby neuron-specific enhancement in excitability could play critical roles in the emergence of cells that encode a specific context^2,10,38–40^.

We postulate that the activity-dependent increase in firing rate demonstrated in our study (**Fig. 2**), along with other forms of plasticity, could assist in the formation of engram cells (**Fig. 10**). It is well established that context-dependent afferent inputs to the dentate gyrus activate a subset of DG granule cells, which elicit action potentials in theta-burst patterns^17,18^. Such sparse recruitment of specific neurons during specific behavioral contexts has been shown to rely on several factors, including targeted connectivity driven by adult neurogenesis, targeted local inhibitory circuits and heterogeneities in the excitability of granule cells^7,39–46^. Our study demonstrates that behaviorally observed theta-burst pattern of action potentials *enhances* the ability of neurons to fire more action potentials, specifically in those neurons that were recruited by afferent activity to elicit such patterns of activity. Such neuron-specific enhancement of excitability in specific neurons that were recruited during a specific context constitutes the core of engram cell formation. We postulate that the form of plasticity reported in this study, along with the underlying cellular and ion channel mechanisms, could provide a mechanistic basis for the emergence of engram cells that encode for a given context (**Fig. 10**).

**Figure 10.**
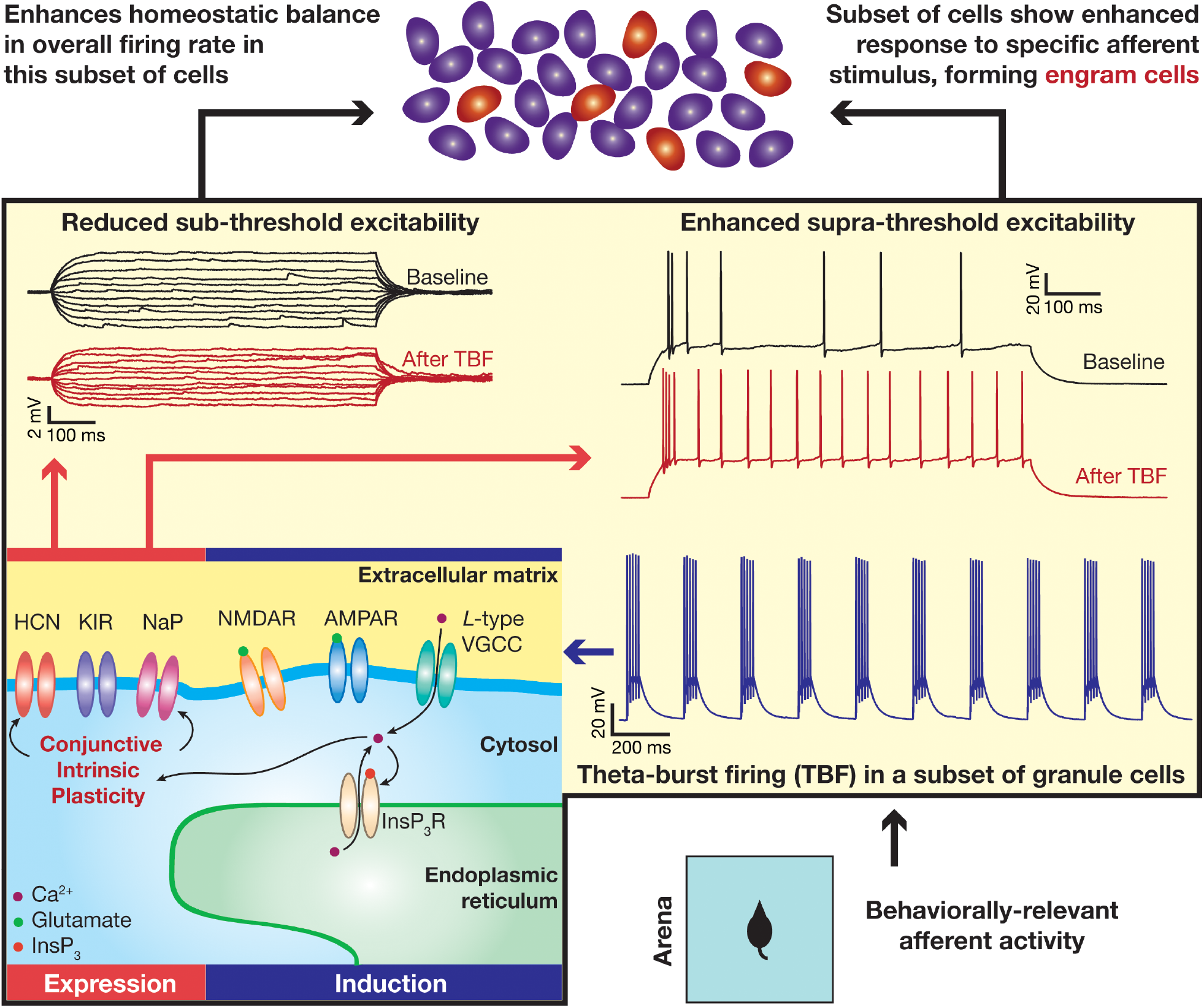
The expression of and implications for activity-dependent plasticity manifolds. Behavioral context-dependent afferent inputs to the dentate gyrus recruit a subset of DG granule cells, which elicit action potentials in theta-burst patterns. Our study shows that theta-burst firing reduces sub-threshold excitability, but contrastingly increases supra-threshold excitability. Mechanistically, *induction* of this form of plasticity is dependent on cytosolic calcium influx through *L*-type voltage-gated calcium channels (VGCC) and inositol trisphosphate receptors (InsP_3_R). The *expression* of TBF-induced contrasting plasticity in sub- and supra-threshold excitability is an outcome of conjunctive plasticity in multiple ion channels: hyperpolarization-activation cyclic-nucleotide-gated (HCN), inward rectifier potassium (KIR) and persistent sodium (NaP) channels. Whereas synergistic interactions between plasticity in HCN and KIR channels mediate the sub-threshold reduction in excitability (represented as a reduction of input resistance), plasticity in NaP channels mediated the enhancement of supra-threshold excitability (represented as enhanced action-potential firing). These findings of the study are shown within the yellow box. The enhanced supra-threshold excitability in the subset of contextually recruited cells (which manifested TBF) could result in the emergence of “engram cells” (highlighted in red in the top panel of GC granule cells), which *encode* for this specific behavioral context. The concomitant reduction in sub-threshold excitability, on the other hand, provides overall *homeostatic balance* in this subset of cells, to counteract the enhanced firing and enable *stable encoding* in a subset of DG granule cells specifically recruited by the behavioral context.

In this context, the manifestation of cell-to-cell variability (**Figs. 2–3, Supplementary Fig. S2**) in activity-dependent intrinsic plasticity could form a cellular substrate for why certain neurons (and not others) are primed to become engram cells^39,40,43,45^. Specifically, across measurements, whereas certain neurons showed tremendous changes in response to the same activity pattern, others did not (**Fig. 2–4; Supplementary Figs. S2**). Additionally, expression of enhancement of supra-threshold excitability as a consequence of TBF was reliant on a *competition* between two opposing changes (in sub-threshold excitability and in AP threshold). Together, the sparse and orthogonal connectivity patterns in the DG, in conjunction with such competition among plasticity in different components could play a critical role in regulating resource allocation for memory storage^40,42–44^.

### Conjunctive plasticity in multiple ion channels points to plasticity manifolds

We demonstrated that the *same* plasticity protocol induces conjunctive plasticity in different channels. We argue that such conjunctive plasticity points to the presence of a tightly regulated *plasticity manifold*, whereby changes to different cellular components are not arbitrary but follow specific rules enforced by common or coupled molecular signaling cascades regulating such plasticity. The plasticity manifold is reminiscent of neuronal intrinsic manifold and behavioral manifold defined in the context of natural *vs.* unnatural neuronal activity and behavior^20,21^. Akin to tightly interconnected patterns of activity across neurons, plasticity in neural components, despite being ubiquitous, seems to follow specific rules on how different components *covary*^3,4,6,47,48^. These rules are enforced by signaling cascades that target specific sets of downstream components, in a manner that seems to also depend on the specific patterns of afferent inputs and the input state of the neuron^4^. Our results show a further instance of the manifestation of such plasticity manifold whereby three distinct channels change concomitantly, introducing conjunctive changes resulting in competition among them to define supra-threshold excitability.

There are advantages of tightly regulated plasticity manifolds involving conjunctive plasticity in different structural components. For instance, our study presents a possible intrinsic route that a neuron could pursue to concomitantly achieve homeostasis while encoding or storage of information (**Fig. 10**). Specifically, from the perspective of engram formation, we had earlier postulated that the activity-dependent increase in firing rate could assist the formation of engram cells that encode for a given context. Within such a postulate, the concomitant reduction in sub-threshold excitability would ensure that the response of the cell to other behavioral contexts is suppressed, apart from maintaining homeostatic balance (to compensate for the enhanced firing rate of the cell). Thus, the *contrasting* patterns of plasticity in sub- *vs.* supra-threshold excitability could be a cell-autonomous substrate to encode new information through enhanced excitability, accompanied by a mechanism to enhance specificity to specific contexts and to enable homeostatic balance of overall afferent drive to the neuron^6^. This is reminiscent of plasticity manifolds in CA1 pyramidal neurons, whereby the *same* theta-burst pairing protocol induces putative *mnemonic* changes in synaptic strength^49^, in transient potassium channels, and in SK channels^50^, apart from concomitantly inducing putative *homeostatic* changes in HCN channels^22,23^. In our study, we show that the *mnemonic* role could be played by plasticity in persistent sodium channels and the *homeostatic* part could be played by changes in HCN and inward-rectifier potassium channels. Thus, conjunctive plasticity involving multiple neural components forms an ideal substrate for effectively achieving the twin goal of encoding and homeostasis^4^ in a cell-type dependent manner, with plasticity manifolds acting as a mechanistic basis for population activity to remain within the neuronal activity manifold. Within this framework, the strong rules governing *concomitant plasticity in multiple components* ensure that the encoding and homeostasis processes don’t interfere with each other, thereby avoiding catastrophic forgetting or unstable learning and enabling continual stable learning.

### Implications for activity-dependent intrinsic plasticity to metaplasticity and channelopathies

It is now well established that plasticity in intrinsic properties could result in metaplasticity through changes in excitability and in temporal summation^51–54^. Our results, demonstrating changes to temporal summation and to sub- and supra-threshold excitability would therefore result in metaplasticity in synaptic plasticity profiles of the dentate gyrus. Such metaplasticity introduced intrinsic plasticity could also form a putative substrate for changes in threshold for synaptic plasticity by *prior* theta-frequency synaptic activity^55,56^. Specifically, prior theta-frequency synaptic activity could have induced changes in intrinsic properties, which in turn could have altered the synaptic plasticity profiles resulting in altered thresholds observed under certain experimental conditions.

From the pathophysiological perspective, altered activity patterns have been associated with intrinsic plasticity in the dentate gyrus, including in channels explored in this study^25,57–62^. Specifically, conjunctive changes in HCN and K_ir_ channels reported in this study is reminiscent of changes in these two channels observed in human DG granule cells with temporal-lobe epilepsy^25^. Therefore, the cellular and ion channel mechanisms associated with intrinsic plasticity explored in this study could play a role in the presence of pathological activity patterns as well. These channelopathies could be homeostatic or could exacerbate the pathological activity patterns in a manner that is dependent on several internal and external factors.

### Limitations, caveats and future directions

Although we do not know the specific reasons for a time-dependent reduction in input resistance in the presence of riluzole (**Fig. 7**), it could be a consequence of the action of riluzole on one of its several known targets. Among these is the ability of riluzole to activate two-pore domain potassium channels, targeting specific subunits^63^ that are known to express and alter excitability of DG granule cells^64,65^. Such activation of two-pore domain potassium channels could potentially explain the time dependent reduction in input resistance we observed.

With reference to the mechanistic basis of TBF-induced plasticity, future studies should more thoroughly explore the different calcium sources participating in the induction of this plasticity. Our study unveils a role for ER calcium in this form of plasticity, and suggests AP-induced influx of calcium through *L*-type calcium channels as a source for initiating such release through InsP_3_ receptors. In addition to these, future studies should explore the roles of different metabotropic receptors, ryanodine receptor and other voltage-gated calcium channels in mediating TBF-induced plasticity^35,36,66–69^. Importantly, future studies should explore the specific signaling cascades, downstream of calcium influx, that mediate the conjunctive plasticity in these ion channels.

Although our focus in this study was on the somata of DG granule cells and on three specific ion channels, it is possible that other channels might change in response to TBF and the manifestation of plasticity might be distinct at different somato-dendritic locations. Therefore, future studies should assess plasticity in other channels employing whole-cell current clamp recordings, voltage imaging, calcium imaging, pharmacological blockade and cell-attached recordings for locations spanning the somato-dendritic axis of DG granule cells. This is important because the localization profile of plasticity is critical in assessing their implications to specific functions, and such analyses would provide the complete span of the plasticity manifold with reference to this form of intrinsic plasticity.

Future studies should also explore TBF-induced intrinsic plasticity across the dorso-ventral, superficial-deep and infrapyramidal-suprapyramidal axes of the dentate gyrus^11^ to assess potential differences in plasticity expression across different subregions of the dentate gyrus. In this context, although there are several studies exploring differences in synaptic plasticity profiles in mature *vs.* immature neurons of the DG^7,8,13^, potential differences in intrinsic plasticity profiles between mature *vs.* immature neurons has not been thoroughly explored. These differences might play a role in how immature neurons integrate into a functional network, and how they participate in engram formation with reference to a specific context. Along the same lines, although the role of different neuromodulators on synaptic plasticity has been explored more thoroughly, the impact of neuromodulatory inputs on intrinsic plasticity profiles has not been explored. Future studies should explore the role of adult neurogenesis and neuromodulators in induction and expression of intrinsic plasticity.

## MATERIALS AND METHODS

### Ethical approval

All experiments reported in this study were performed in strict adherence to the protocols cleared by the Institute Animal Ethics Committee (IAEC) of the Indian Institute of Science, Bangalore. Animals were provided *ad libitum* food and water and were housed with an automated 12 h light– 12h dark cycle. All animals were obtained from the in-house breeding setup at the central animal facility of the Indian Institute of Science. Surgical and electrophysiological procedures were similar to previously established protocols^23,26,46,70^ and are detailed below.

### Slice preparation for *in-vitro* patch clamp recording

Male Sprague-Dawley rats of 6- to 8-weeks age were anesthetized by intraperitoneal injection of a ketamine-xylazine mixture. After onset of deep anesthesia, assessed by cessation of toe-pinch reflex, transcardial perfusion of ice-cold cutting solution was performed. The cutting solution contained 2.5 mM KCl, 1.25 mM NaH_2_PO_4_, 25 mM NaHCO_3_, 0.5 mM CaCl_2_, 7 mM MgCl_2_, 7 mM dextrose, 3 mM sodium pyruvate, and 200 mM sucrose (pH 7.3, ~300 mOsm) saturated with 95% O_2_ and 5% CO_2_. Thereafter, the brain was removed quickly and 350-μm thick near-horizontal slices were prepared from middle hippocampi (Bregma, –6.5 mm to –5.1 mm), using a vibrating blade microtome (Leica Vibratome), while submerged in ice-cold cutting solution saturated with 95% O_2_ and 5% CO_2_. The slices were then incubated for 10–15 mins at 34° C in a chamber containing the holding solution (pH 7.3, ~300 mOsm) with the composition of: 125 mM NaCl, 2.5 mM KCl, 1.25 mM NaH_2_PO_4_, 25 mM NaHCO_3_, 2 mM CaCl_2_, 2 mM MgCl_2_, 10 mM dextrose, 3 mM sodium pyruvate saturated with 95% O_2_ and 5% CO_2_. The slices were kept in a holding chamber at room temperature for at least 45 min before the start of recordings.

### Electrophysiology: Whole-cell current-clamp recording

For electrophysiological recordings, slices were transferred to the recording chamber and continuously perfused with carbogenated artificial cerebrospinal fluid (ACSF/extracellular recording solution) at a flow rate of 2–3 mL/min. All neuronal recordings were performed under current-clamp configuration at physiological temperatures (32–35° C), achieved through an inline heater that was part of a closed-loop temperature control system (Harvard Apparatus). The carbogenated ACSF contained 125 mM NaCl, 3 mM KCl, 1.25 mM NaH_2_PO_4_, 25 mM NaHCO_3_, 2 mM CaCl_2_, 1 mM MgCl_2_, 10 mM dextrose (pH 7.3; ~300 mOsm). Slices were first visualized under a 10× objective lens to locate the granule cell layer in the crest sector of the dentate gyrus. Then, a 63× water immersion objective lens was employed to perform patch-clamp recordings from DG granule cells in the crest sector, through a Dodt contrast microscope (Carl Zeiss Axioexaminer). Whole-cell current-clamp recordings were performed from visually identified dentate gyrus granule cell somata, using Dagan BVC-700A amplifiers. Electrophysiological signals were low-pass filtered at 5 kHz and sampled at 10–40 kHz. All data acquisition and analyses were performed using custom-written software in Igor Pro (Wavemetrics).

Images of cell location were captured for record and for post-facto confirmation of the region where the cell belonged. Borosilicate glass electrodes with resistance between 2–6 MΩ (more often electrodes with ~4 MΩ electrode resistance were used) were pulled (P-97 Flaming/Brown micropipette puller; Sutter) from thick glass capillaries (1.5 mm outer diameter and 0.86 mm inner diameter; Sutter) and used for patch-clamp recordings. The pipette solution contained 120 mM K-gluconate, 20 mM KCl, 10 mM HEPES, 4 mM NaCl, 4 mM Mg-ATP, 0.3 mM Na-GTP, and 7 mM K2-phosphocreatine (pH 7.3 adjusted with KOH, osmolarity ~300 mOsm).

Series resistance was monitored (once every 30 s) and compensated online using the bridge-balance circuit of the amplifier. Experiments were discarded only if the initial resting membrane potential was more depolarized than −60 mV or if series resistance rose above 30 MΩ, or if there were fluctuations in temperature and ACSF flow rate during the course of the experiment. Unless otherwise stated, experiments were performed at the initial resting membrane potential (reported here as *V*_RMP_) of the cell. Voltages have not been corrected for the liquid junction potential, which was experimentally measured to be ~8 mV.

### Sub-threshold measurements

We characterized plasticity in intrinsic properties of DG granule neurons using several electrophysiological measurements obtained through several pulse-current and frequency-dependent current injections^9,23,26,70^. Input resistance (*R*_in_) was measured as the slope of a linear fit to the steady-state *V-I* plot obtained by injecting subthreshold current pulses of amplitudes spanning −25 to +25 pA, in steps of 5 pA (**Fig. 1d**). To assess temporal summation, five α-excitatory postsynaptic potentials (α-EPSPs) with 50 ms interval were evoked by current injections of the form *I*_α_ =*I*_max_*t* exp (–α*t*), with α = 0.1 ms^−1^ (**Fig. 1g**). Temporal summation ratio (*S*_α_) in this train of five EPSPs was computed as *E*_last_/*E*_first_, where *E*_last_ and *E*_first_ were the amplitudes of last and first EPSPs in the train, respectively. Percentage sag was measured from the voltage response of the cell to a hyperpolarizing current pulse of 100 pA (**Fig. 1b**, top) and was defined as 100 (1–*V*_ss_/*V*_peak_), where *V*_ss_ and *V*_peak_ depicted the steady-state and peak voltage deflection from *V*_RMP_, respectively.

The chirp stimulus (**Fig. 1b**) used for characterizing the impedance amplitude (ZAP) profiles was a sinusoidal current of constant amplitude below firing threshold, with its frequency linearly spanning 0–15 Hz in 15 s (*Chirp15*). The magnitude of the ratio of the Fourier transform of the voltage response (**Fig. 1e**) to the Fourier transform of the *Chirp15* stimulus formed the ZAP (**Fig. 1f**):

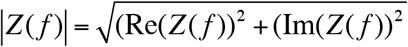

where Re(*Z*(*f*)) and Im(*Z*(*f*)) refer to the real and imaginary parts of the impedance *Z* as a function of frequency *f*. The maximum value of impedance across all frequencies was measured as the maximum impedance amplitude (|*Z*|_max_; **Fig. 1f**). The frequency at which the impedance amplitude reached its maximum was the resonance frequency (*f*_R_). Resonance strength (*Q*) was measured as the ratio of the maximum impedance amplitude to the impedance amplitude at 0.5 Hz^23^.

### Supra-threshold measurements

To understand the possible cellular mechanisms underlying the expression of activity-dependent intrinsic plasticity, we further analyzed the various features of action potentials. First, AP firing frequency was computed by extrapolating the number of spikes obtained during a 700 ms current injection to 1 s. Current amplitude of these pulse-current injections was varied from 0 pA to 250 pA in steps of 50 pA, to construct the firing frequency *vs.* injected current (*f–I*) plot (**Fig. 1h–i**). Various AP related measurements^9^ were derived from the voltage response of the cell to a 250 pA pulse-current injection. AP amplitude (*V*_AP_) was computed as the difference between the peak voltage of the spike 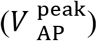 and *V*_RMP_. The temporal distance between the timing of the first spike and the time of current injection was defined as latency to first spike (*T*_1AP_). The duration between the first and the second spikes was defined as the first inter-spike interval (*T*_1ISI_). AP half-width (*T*_APHW_) was the temporal width measured at the half-maximal points of the AP peak with reference to *V*_RMP_. The 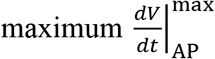, and 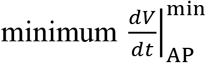 values of the AP temporal derivative were calculated from the temporal derivative of the AP trace. The voltage in the AP trace corresponding to the time point at which the d*V*/d*t* crossed 20 V/s defined AP threshold. All supra-threshold measurements were obtained through current injections into the cell resting at *V*_RMP_.

### Theta Burst Firing Protocol

Experimental procedures for inducing theta-burst firing (TBF) were similar to previously established protocols^22,23^. To induce TBF, we employed 3 trains of 10 theta-modulated bursts each, separated by 10 s. Each train (**Fig. 1c**) was made of a burst of 5 APs with an intra-burst frequency of 100 Hz (10 ms ISI), and the inter-burst frequency set at 5 Hz (200 ms). Each AP within the burst was initiated by injecting a large current (2 nA) of small duration (2 ms) into the neuron. The experimental protocol (**Fig. 1b–c**) for TBF involved initial measurements of the *V-I* curve, *f-I* curve and *S*_α_, followed by establishment of a 5 min stable baseline while monitoring resting membrane potential and *R*_in_. Monitoring of these measurements were from cell responses to the *Chirp15* stimulus, measured twice every minute, during this initial 5 min period and for 40 min after TBF, followed by a final measurement of the *V-I* curve, *f-I* curve and *S*_α_. We recorded from one selected cell in any given slice, as the induction protocol could have had unknown effects on neighboring cells through synaptic transmission or other signaling cascades.

To assess pairwise relationship across 13 different sub- and supra-threshold measurements, we analyzed the scatter plot matrices of post-TBF changes in these measurements (**Fig. 3a**). We computed Pearson’s correlation coefficients for each of these pair-wise scatter plots and analyzed the distribution of correlation coefficients (**Fig. 3**).

### Pharmacological Blockers

#### Synaptic receptor blockers

Drugs and their concentrations used in the experiments were 10 μM 6-cyano- 7-nitroquinoxaline-2,3-dione (CNQX), an AMPA receptor blocker; 10 μM (+) bicuculline and 10 μM picrotoxin, both GABA_A_ receptor blockers; 50 μM d,l-2-amino-5-phosphonovaleric acid (d,l-APV), NMDA receptor and 2 μM CGP55845, GABA_B_ blocker (all synaptic blockers from Allied Scientific) in the bath solution. To block *Inositol trisphosphate (InsP_3_) receptors*, 1 mg/mL heparin (20,000–25,000 molecular weight; Calbiochem) was included in the recording pipette.

#### Voltage-gated ion channel blockers

20 μM ZD7288 (Allied Scientific) or 20 μM zatebradine (Sigma Aldrich) was employed to block HCN channels. 50 μM BaCl_2_ (Sigma Aldrich) or 0.3 μM tertiapin-Q (Tocris Biosciences) was used to block inward-rectifier potassium channels. 20 μM Riluzole (Allied Scientific) and 10 μM Nimodipine (Tocris Biosciences) were used to block persistent sodium and *L*-type calcium channels, respectively. For experiments with ZD7288, cells were patched with pipette solution containing 20 μM ZD7288 along with adding it to bath solution^70^. For long-term control and TBF experiments in the presence of pharmacological agents, slices were pretreated with the respective pharmacological agent for at least 15 min before the start of recordings.

#### Fast calcium chelator

30 mM (1,2-bis(o-aminophenoxy)ethane-N,N,N′,N′-tetraacetic acid), BAPTA (Thermo Fisher) was incorporated into the pipette solution. The constituents of the pipette solution were (in mM): 30 mM K_4_-BAPTA, 20 mM KCl, 10 mM HEPES, 4 mM NaCl, 4 mM Mg-ATP, 0.3 mM Na-GTP, and 7 mM K_2_-phosphocreatine (pH 7.3 adjusted with KOH, osmolarity ~300 mOsm adjusted with sucrose).

### Statistics and Reproducibility

All statistical analyses were performed using the R computing package (http://www.r-project.org/). In order to avoid false interpretations and to emphasize the heterogeneities, the entire range of measurements are reported in figures rather than providing only the summary statistics^4^. Necessary care was taken and appropriate controls were performed for each of the drugs used to ensure that there were no time-dependent changes initiated by just the presence of the drug in the bath or in the pipette solution. For all cases, we performed long-term control experiments (no protocol) and TBF experiments in the presence of same quantity of drugs, and report the entire span of measurements corresponding to the outcomes for both sets of experiments. Statistical comparison was performed with their respective long-term controls. Across figures, the statistics employed for data presentation was consistent with the statistical test used to compare two populations of data. Specifically, when data is reported as mean ± SEM, parametric tests (paired or unpaired Student’s *t* test) were employed, and when data is reported as median (along with the entire distribution of the data or the quartiles), we employed non-parametric tests (Wilcoxon ranked sum or signed rank tests). Results of statistical tests, with exact *p* values and the name of the statistical test employed, are provided in the figure panels or in the respective figure legends.

## ACKNOWLEDGMENTS

The authors thank Dr. Daniel Johnston, Dr. Sufyan Ashhad, Dr. Richard Gray, Dr. M. K. Mathew and the members of the cellular neurophysiology laboratory for helpful discussions and/or for comments on a draft of this manuscript. This work was supported by the Wellcome Trust-DBT India Alliance (Senior fellowship to RN; IA/S/16/2/502727), Human Frontier Science Program (HFSP) Organization (RN), the Department of Biotechnology through the DBT-IISc partnership program (RN), the Revati & Satya Nadham Atluri Chair at IISc (RN), the Department of Science and Technology (RN), and the Ministry of Human Resource Development (RN & PM).

## AUTHOR CONTRIBUTIONS

P. M. and R. N. designed experiments; P. M. performed experiments and carried out data analysis; P. M. and R. N. co-wrote the paper.

