## Supplementary Figures S1-S13; Supplementary Table S1 for "Plasticity manifolds: Conjunctive changes in multiple ion channels mediate activity-dependent plasticity in hippocampal granule cells"

### <u>Supplementary Material</u>

Supplementary Figure S1 .......... 2
Supplementary Figure S2 .......... 3
Supplementary Figure S3 .......... 4
Supplementary Figure S4 .......... 5
Supplementary Figure S5 .......... 6
Supplementary Figure S6 .......... 7
Supplementary Figure S7 .......... 8
Supplementary Figure S8 .......... 9
Supplementary Figure S9 .......... 10
Supplementary Figure S10 .......... 11
Supplementary Figure S11 .......... 12
Supplementary Figure S12 .......... 13
Supplementary Figure S13 .......... 14

Supplementary Table S1 .......... 15

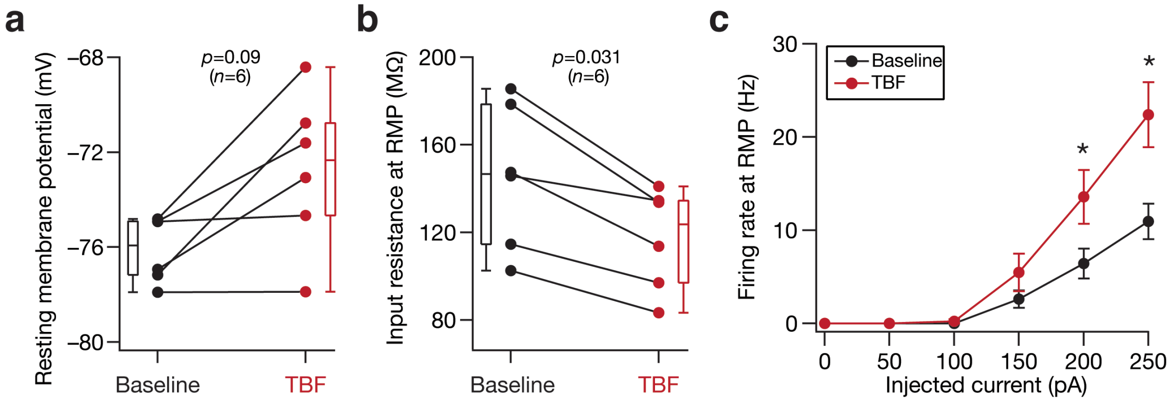


**Supplementary Figure S1. Contrasting plasticity in sub- *vs.* supra-threshold excitability was observed when measurements were recorded at respective resting membrane potentials.** (**a**) Population data of resting membrane potential from all DG granule cells recorded before (black) and 40 mins after (red) TBF protocol, showing persistent changes in RMP after TBF. (**b–c**) Population data of measurements from all DG granule cells recorded before (black) and 40 mins after (red) TBF protocol. Measurements were performed at the respective resting membrane potentials (shown in panel **a** for each cell), with zero current injection. Input resistance obtained from the *V–I* plot reduced after TBF (**b**), whereas firing rate increased (**c**). The Wilcoxon signed rank test was employed for *p* value calculation in panels **a–b**, and Student's *t* test was employed for *p* value calculation in panel (**c**). *: $p<0.05$.

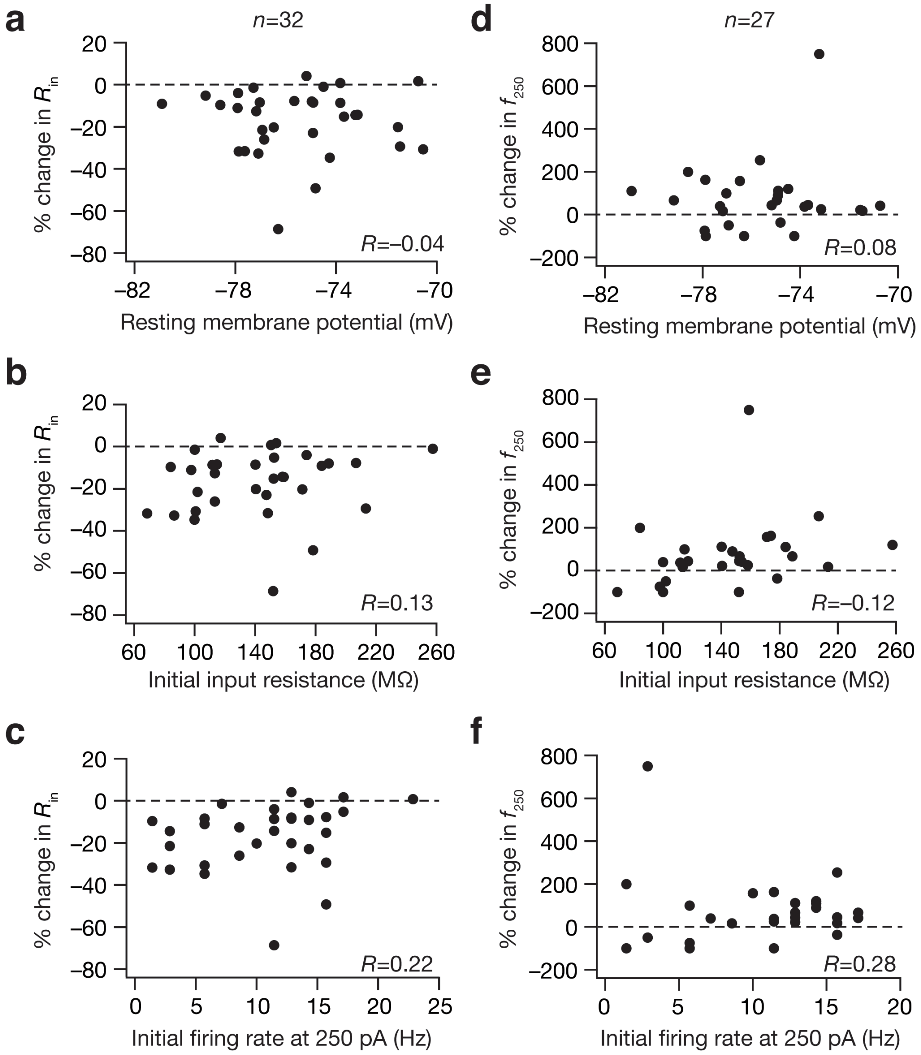


**Supplementary Figure S2. Weak correlations between heterogeneities in TBF-induced plasticity and heterogeneities in intrinsic properties of DG granule cells.** (**a–c**) Population data representing TBF-induced change in input resistance plotted against resting membrane potential of the cells (**a**); input resistance of the cells (**b**); and firing rate of the cells in response to 250 pA current injection (**c**). (**d–f**) Same as (**a–c**), but with TBF-induced change in firing rate at 250 pA on the ordinate. Pearson correlation coefficient (*R*) is provided for each panel.

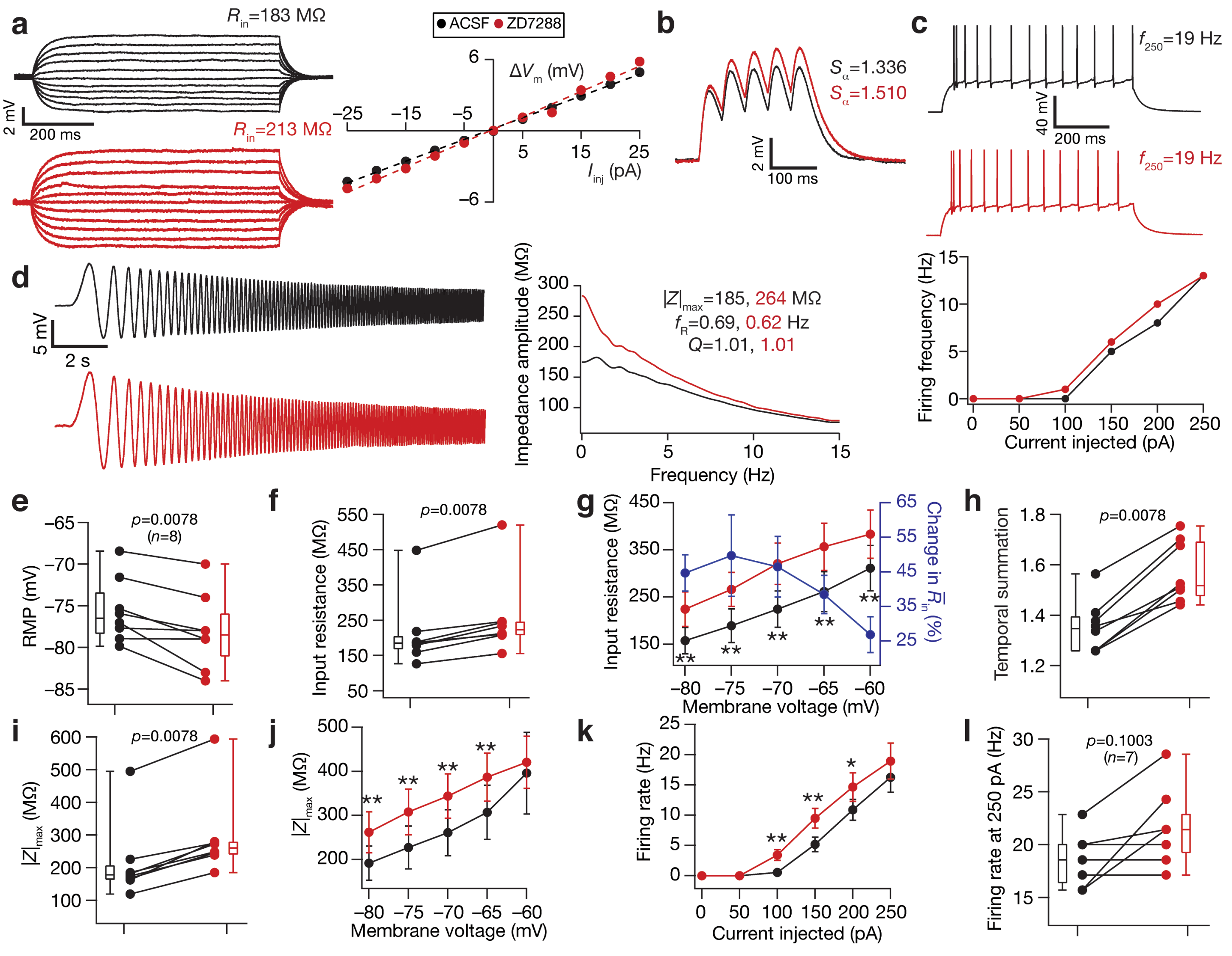


**Supplementary Figure S3. Acute treatment with ZD7288, a HCN channel blocker, hyperpolarized resting membrane potential and enhanced sub- and supra-threshold excitability of DG granule cells.** (**a**) *Right*, Voltage responses of a DG granule cell to 700 ms current pulses of amplitude varying from –25 pA to +25 pA (in steps of 5 pA), with normal ACSF (*black*) and in the presence of 20 µM ZD7288 in the bath (*red*). *Left*, Input resistance ($R_{in}$) was calculated as the slope of the *V–I* plot depicting steady-state voltage response as a function of the injected current amplitude. (**b**) Voltage response of the example neuron to 5 alpha-current injections arriving at 20 Hz, depicting temporal summation. Temporal summation ratio ($S_\alpha$) was computed as the ratio of the amplitude of the fifth response to that of the first. (**c**) *Top,* Voltage response of the example neuron to a 700-ms current pulse of 250 pA in ACSF (*black*) and in the presence of ZD7288 (*red*). *Bottom*, Frequency of firing plotted as a function of injected current amplitude for the example cell. Note that these are firing frequencies converted from the number of spikes obtained for 700-ms duration pulses. (**d**) *Left*, Voltage responses of the example neuron to the chirp current (**Fig. 1b**, top) in ACSF (*black*) and in the presence of ZD7288 (*red*). *Right*, Impedance amplitude computed from the current stimulus shown in **Fig. 1b** (*top*) and the voltage responses shown on the left. $|Z|_{max}$ represents the maximum impedance amplitude, $Q$ is resonance strength and resonance frequency is represented by $f_R$. (**e–l**) Population data of measurements from all DG granule cells recorded before and after adding ZD7288 to the bath: RMP (**e**); input resistance, $R_{in}$ (**f**); membrane potential dependence of input resistance (***g***); temporal summation (**h**); impedance amplitude, $|Z|_{max}$ (**i**) and its voltage dependence (**j**); firing rate at 0–250 pA current injection (**k**) and for 250 pA current injection (**l**). The Wilcoxon signed rank test was used for $p$ value calculation in panels **e-f**, **h-i** and l, for comparing measurements from the same set of cells. For panel **g**, **j** and **k** statistical comparisons were performed with paired Student's $t$ test; *: $p < 0.05$, **: $p < 0.005$. All experiments reported in this figure were performed in the presence of 10 µM CNQX, 10 µM (+) bicuculline and 10 µM picrotoxin.

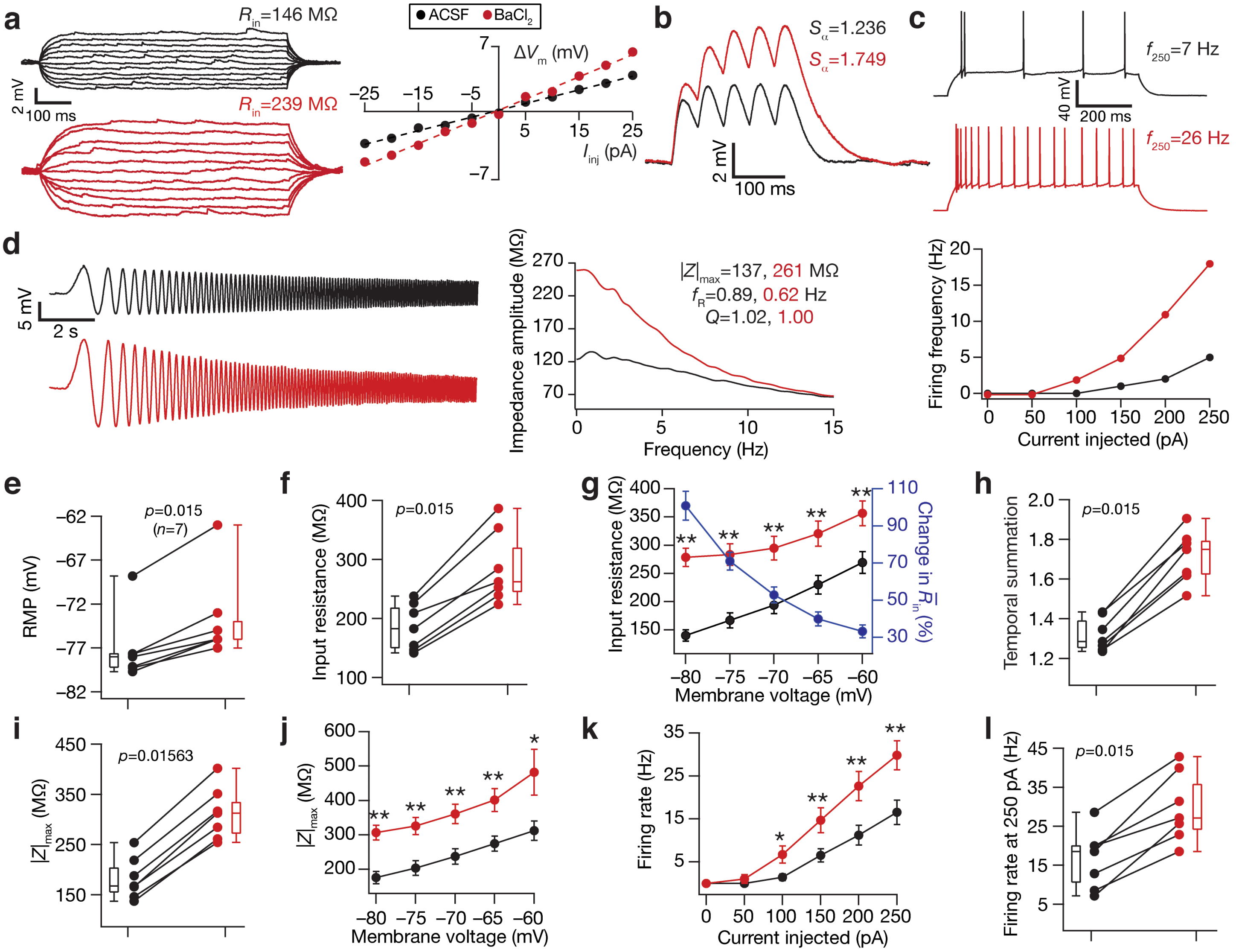


**Supplementary Figure S4. Acute treatment with barium chloride ($BaCl_2$), an inward rectifier potassium channel blocker, depolarized resting membrane potential and enhanced sub- and supra-threshold excitability of DG granule cells.** (**a**) *Right*, Voltage responses of a DG granule cell to 700 ms current pulses of amplitude varying from –25 pA to +25 pA (in steps of 5 pA), with normal ACSF (*black*) and in the presence of 50 µM $BaCl_2$ in the bath (*red*). *Left*, Input resistance ($R_{in}$) was calculated as the slope of the *V–I* plot depicting steady-state voltage response as a function of the injected current amplitude. (**b**) Voltage response of the example neuron to 5 alpha-current injections arriving at 20 Hz, depicting temporal summation. Temporal summation ratio ($S_\alpha$) was computed as the ratio of the amplitude of the fifth response to that of the first. (**c**) *Top,* Voltage response of the example neuron to a 700-ms current pulse of 250 pA in ACSF (*black*) and in the presence of $BaCl_2$ (*red*). *Bottom*, Frequency of firing plotted as a function of injected current amplitude for the example cell. Note that these are firing frequencies converted from the number of spikes obtained for 700-ms duration pulses. (**d**) *Left*, Voltage responses of the example neuron to the chirp current (**Fig. 1b**, top) in ACSF (*black*) and in the presence of $BaCl_2$ (*red*). *Right*, Impedance amplitude computed from the current stimulus shown in **Fig. 1b** (*top*) and the voltage responses shown on the left. $|Z|_{max}$ represents the maximum impedance amplitude, $Q$ is resonance strength and resonance frequency is represented by $f_R$. (**e–l**) Population data of measurements from all DG granule cells recorded before and after adding $BaCl_2$to the bath: RMP (**e**); input resistance, $R_{in}$ (**f**); membrane potential dependence of input resistance (***g***); temporal summation (**h**); impedance amplitude, $|Z|_{max}$ (**i**) and its voltage dependence (**j**); firing rate at 0–250 pA current injection (**k**) and for 250 pA current injection (**l**). The Wilcoxon signed rank test was used for $p$ value calculation in panels **e-f**, **h-i** and **l**, for comparing measurements from the same set of cells. For panel **g**, **j** and **k** statistical comparisons were performed with paired Student's $t$ test; *: $p < 0.05$, **: $p < 0.005$.

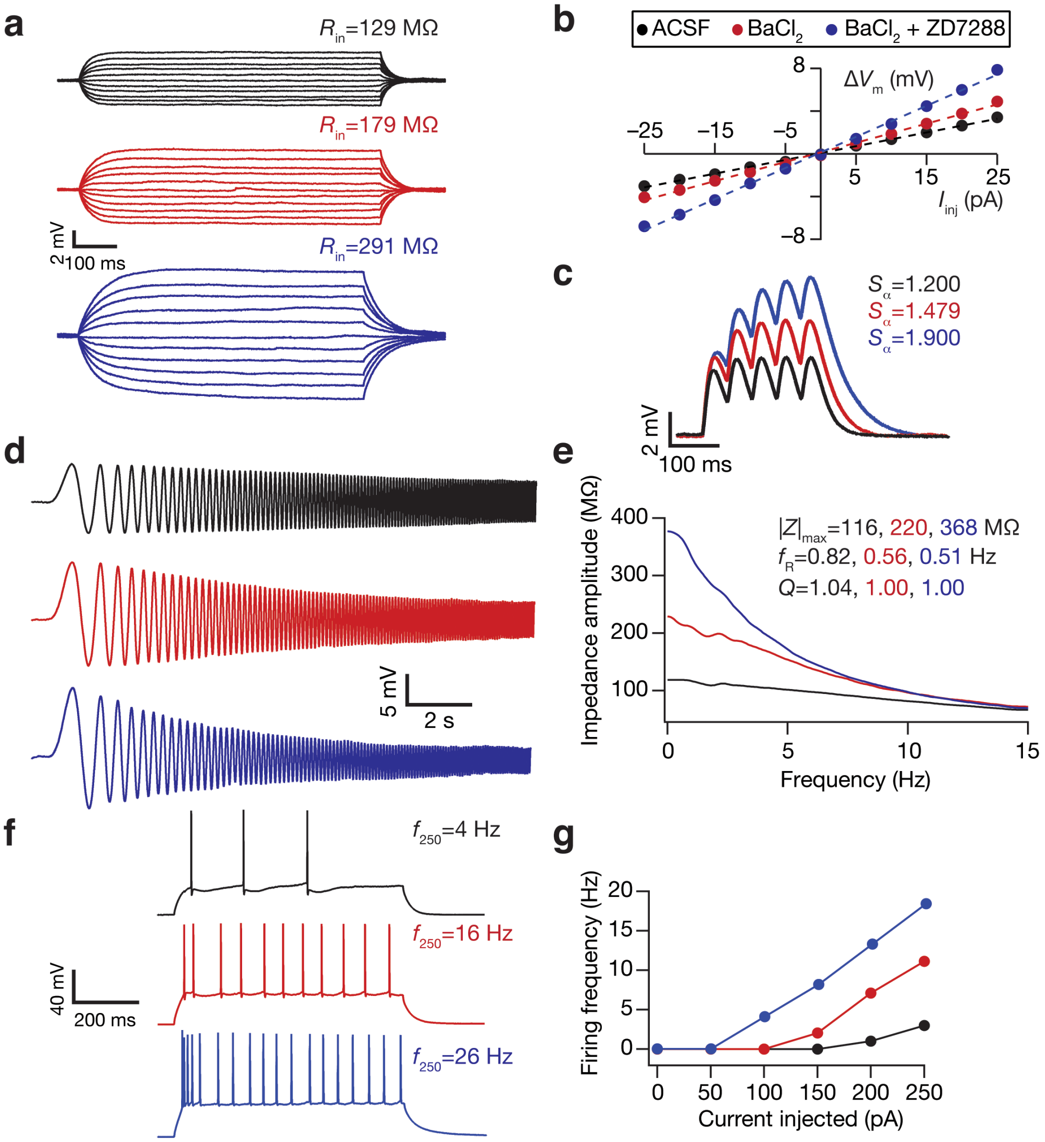


**Supplementary Figure S5. Impact of acute treatment with $BaCl_2$ followed by $BaCl_2$ + ZD7288 on sub- and supra-threshold excitability in an example DG granule cell.** (**a**) Voltage responses of a DG granule cell to 700 ms current pulses of amplitude varying from –25 pA to +25 pA (in steps of 5 pA), with normal ACSF (*black*) and in the presence of 50 μM $BaCl_2$ in the bath (*red*) and in the presence of 50 μM $BaCl_2$ and 20 μM ZD7288 in bath (*blue*). **(b)** Input resistance ($R_{in}$) was calculated as the slope of the *V–I* plot depicting steady-state voltage response as a function of the injected current amplitude. (**c**) Voltage response of the example neuron to 5 alpha-current injections arriving at 20 Hz, depicting temporal summation. Temporal summation ratio ($S_\alpha$) was computed as the ratio of the amplitude of the fifth response to that of the first. (**d**) Voltage responses of the example neuron to the chirp current (**Fig. 1b**, top) in ACSF (*black*), in the presence of $BaCl_2$ (*red*) and in presence of ZD7288 (*blue*). **(e)** Impedance amplitude computed from the current stimulus shown in **Fig. 1b** (*top*) and the voltage responses shown on the left. $|Z|_{max}$ represents the maximum impedance amplitude, $Q$ is resonance strength and resonance frequency is represented by $f_R$. (**f**) *Top,* Voltage response of the example neuron to a 700-ms current pulse of 250 pA in ACSF (*black*), in the presence of $BaCl_2$ (*red*) and in presence of ZD7288 (*blue*). **(g)** Frequency of firing plotted as a function of injected current amplitude for the example cell. Note that these are firing frequencies converted from the number of spikes obtained for 700-ms duration pulses. The experiment reported in this figure was performed in the presence of 10 μM CNQX, 10 μM (+) bicuculline and 10 μM picrotoxin.

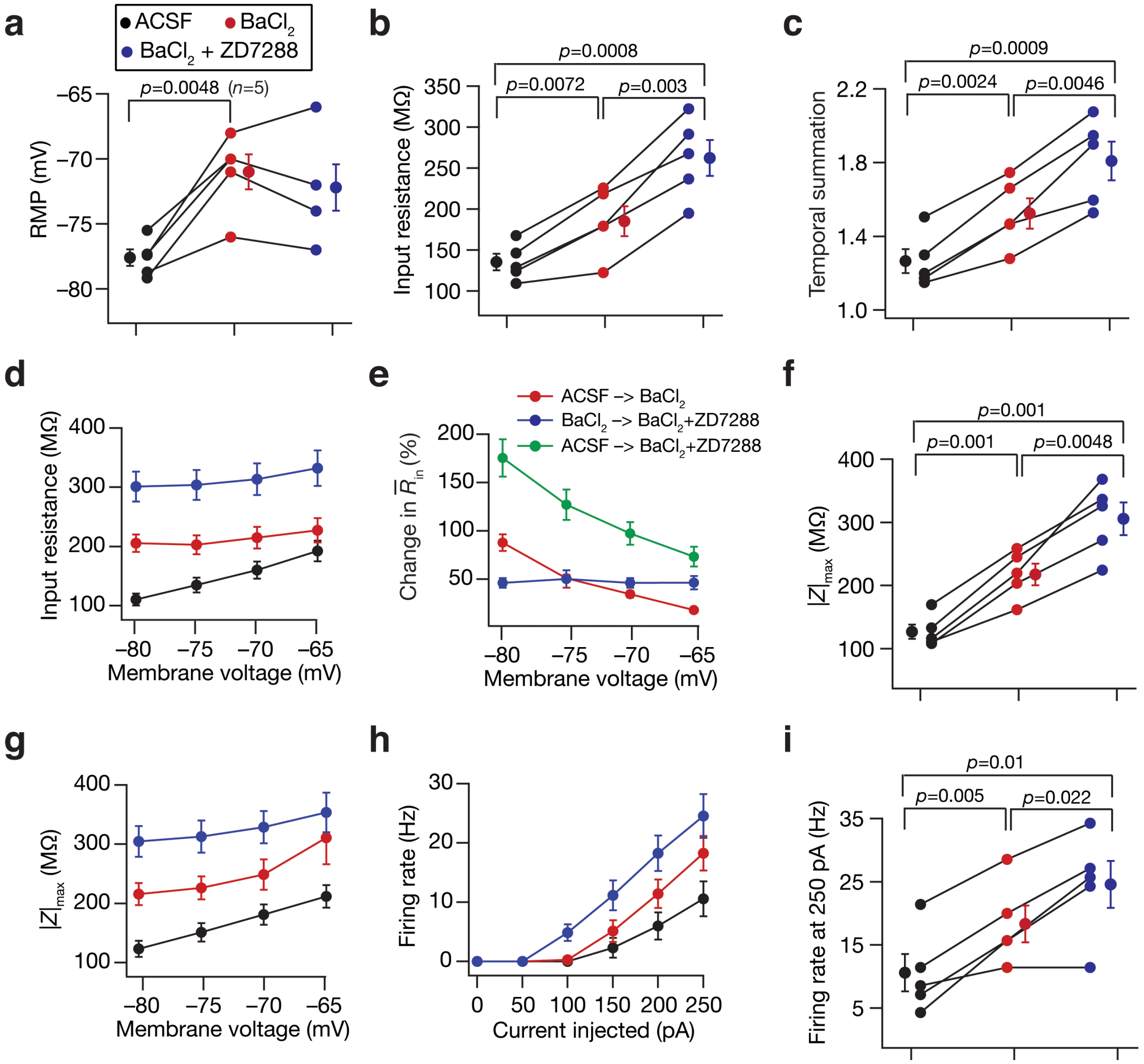


**Supplementary Figure S6. ZD7288 enhanced sub- and supra-threshold excitability of DG granule cells beyond the excitability enhancement induced by $BaCl_2$ treatment.** (**a–i**) Population data of measurements from all DG granule cells recorded in ACSF, then in presence of 50 µM $BaCl_2$ (red) and in the presence of 50 µM $BaCl_2$ and 20 µM ZD7288 in the bath (blue): RMP (**a**); input resistance, $R_{in}$ (**b**); temporal summation (**c**); membrane potential dependence of input resistance (**d**); percentage change in input resistance as a function of membrane potential (**e**): impedance amplitude, $|Z|_{max}$ (**f**) and its voltage dependence (**g**); firing rate for 0–250 pA current injection (**h**) and for 250 pA current injection (**i**). For all panels statistical comparisons were performed with paired Student's *t* test; *: $p < 0.05$, **: $p < 0.005$. For panels **d** and **g**, across-group measurements were significantly different ($p < 0.05$) from each other for all measured voltages. For panel **h**, across-group measurements of action potential firing rates were significantly different ($p < 0.05$) for current injections in the range 150–250 pA; for 100-pA current injection, firing rate in the ($BaCl_2$+ZD7288) group was significantly different ($p < 0.05$) from the other two groups. All experiments reported in this figure were performed in the presence of 10 µM CNQX, 10 µM (+) bicuculline and 10 µM picrotoxin.

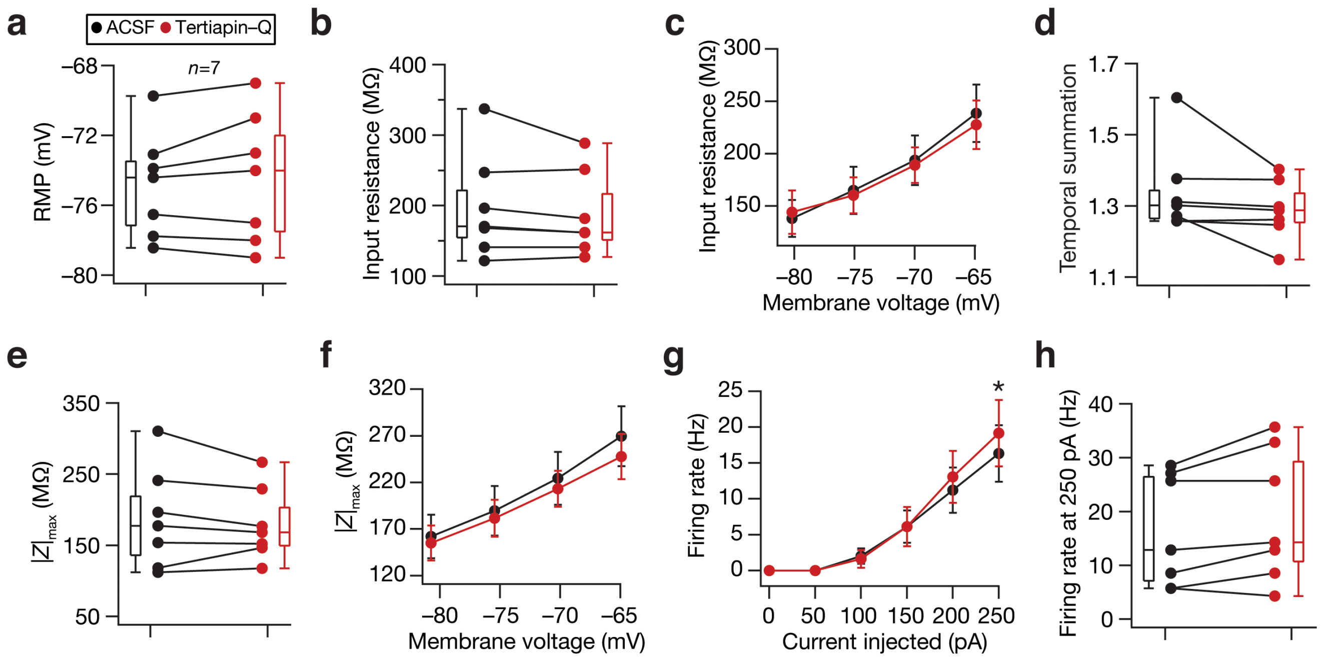


**Supplementary Figure S7. Acute treatment with tertiapin-Q, a blocker of specific subtypes of inward-rectifier potassium channels, yielded no significant change in physiological measurements of DG granule cells.** (**a–h**) Population data of measurements from all DG granule cells recorded before (*black*) and after adding 0.3 µM tertiapin-Q (*red*) to the bath: RMP (**a**); input resistance, $R_{in}$ (**b**); membrane potential dependence of input resistance (*c*); temporal summation (**d**); impedance amplitude, $|Z|_{max}$ (**e**) and its voltage dependence (**f**); firing rate at 0–250 pA current injection (**g**) and for 250 pA current injection (**h**). The Wilcoxon signed rank test was used for $p$ value calculation in panels **a-b, d-e** and **h**, for comparing measurements from the same set of cells. For panel **c**, **f** and **g** statistical comparisons were performed with paired Student's $t$ test; *: $p < 0.05$. All experiments reported in this figure were performed in the presence of 10 µM CNQX, 10 µM (+) bicuculline and 10 µM picrotoxin.

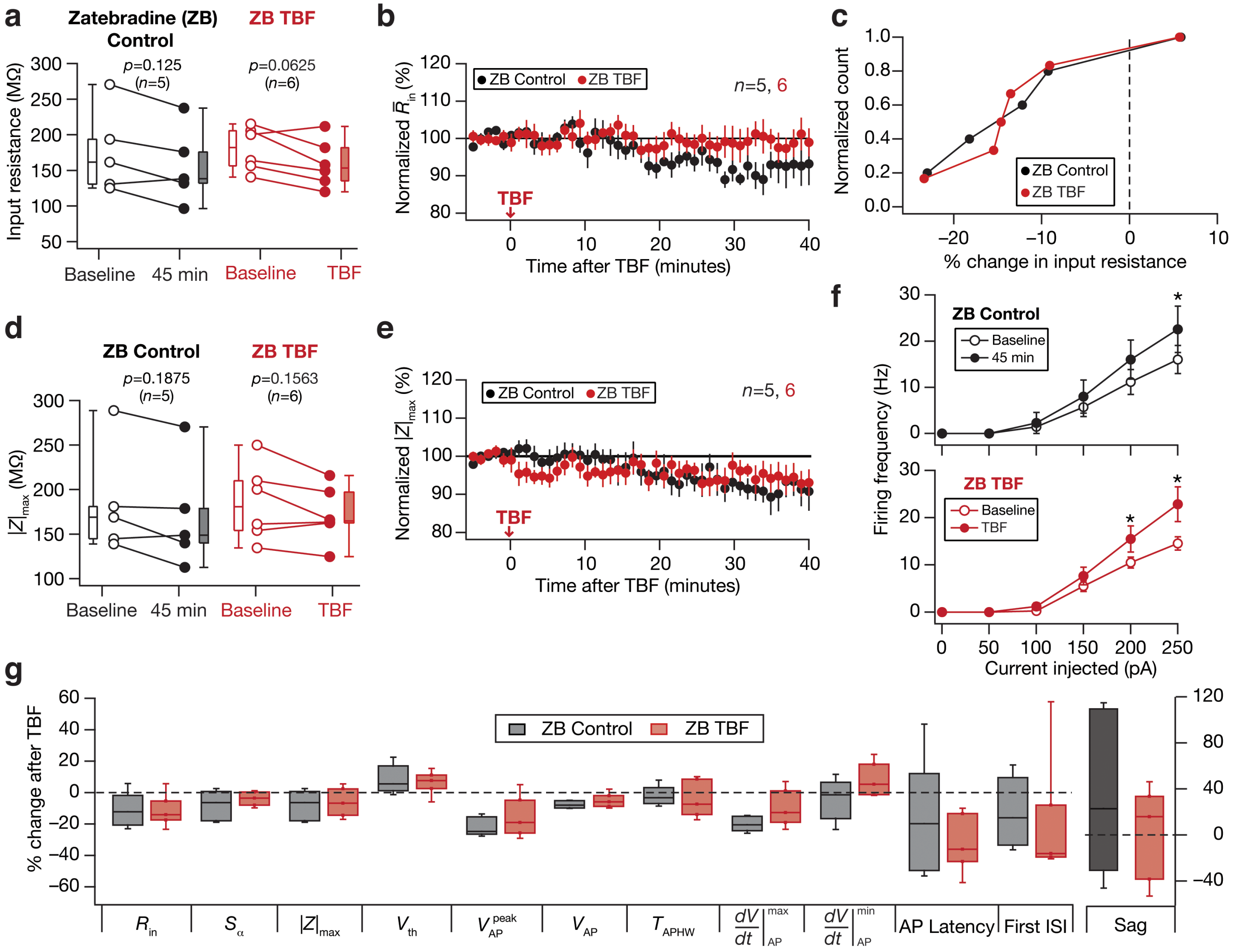


**Supplementary Figure S8. Activity-dependent reduction in sub-threshold excitability was blocked in the presence of 20 µM zatebradine (ZB), a HCN channel blocker.** For all panels, the ZB Control (black) group ($n$=5) corresponds to experiments where no protocol was applied through the 45-min period of the experiment, and ZB TBF (red) group ($n$=5) is for neurons subjected to theta burst firing. For the ZB TBF group, "Baseline" measurements were obtained before TBF and "TBF" measurements were after TBF. All experiments reported in this figure were performed in the presence of 20 µM ZB in the bath. (**a**) Population data representing change in input resistance ($R_{in}$) at the beginning (*empty circles*) and the end (*filled circles*) of the experiment. (**b**) Temporal evolution of percentage change in input resistance estimate ($\bar{R}_{in}$; Mean ± SEM). (**c**) Normalized count of neurons from panel **a** plotted as functions of percentage change in $R_{in}$. (**d–e**) Same as **a–b**, representing impedance amplitude ($|Z|_{max}$). (**f**) Frequency of firing plotted as a function of injected current amplitude for both groups. *: $p$<0.05; **: $p$<0.005. Student's $t$ test. (**g**) Plots comparing percentage change in various measurements from their initial values to end of experiment values. The list of symbols and corresponding measurements is enumerated in Supplementary Table S1. The Wilcoxon signed rank test was used for $p$ value calculation in panels **a** and **d**, for comparing measurements from the same set of cells. The Wilcoxon rank sum test was employed for $p$ value calculation in panel **g**, to compare percentage changes in the control group *vs*. those in the TBF group.

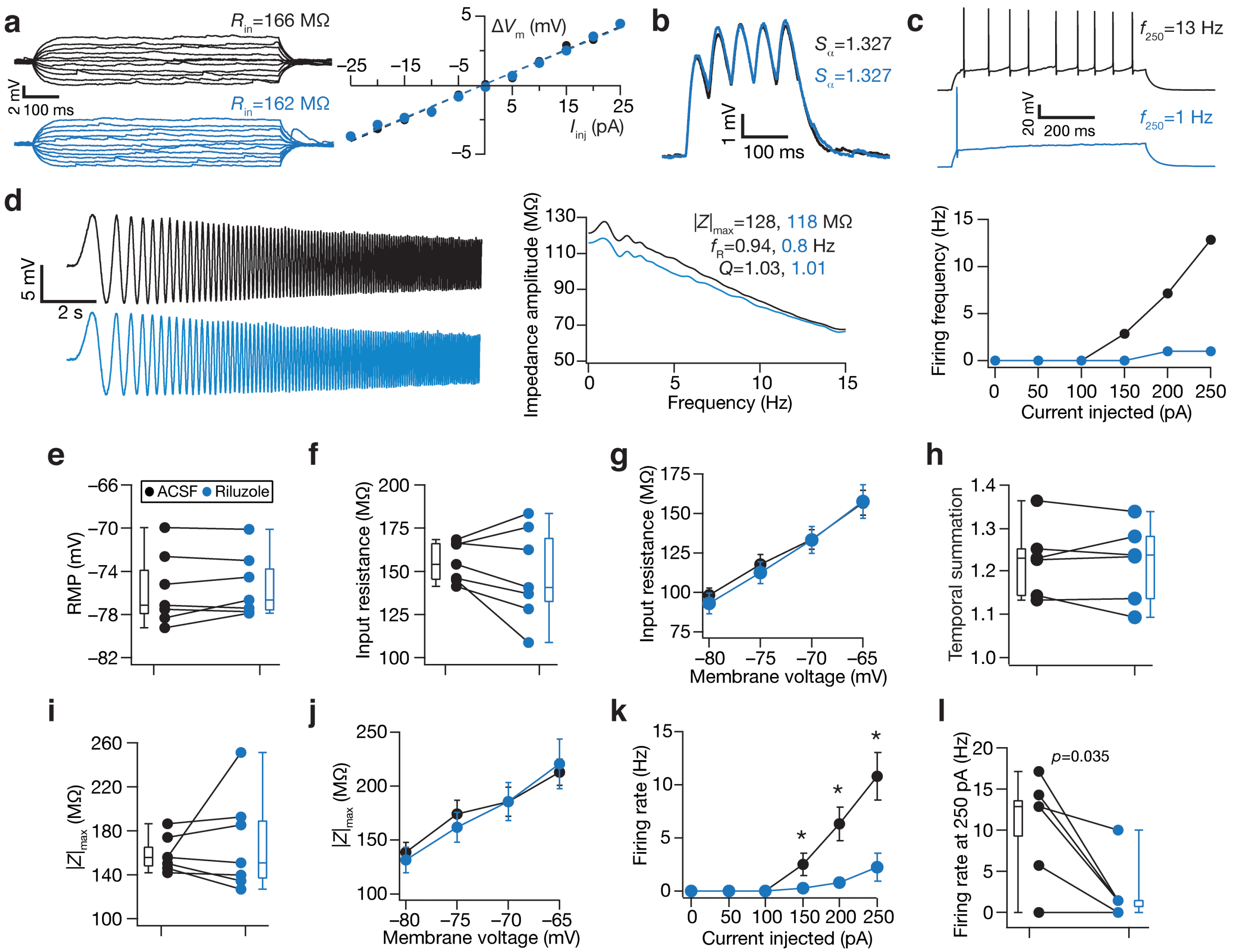


**Supplementary Figure S9. Acute treatment with Riluzole, a persistent sodium channel blocker, reduced the firing rate of DG granule cells without altering their sub-threshold physiological characteristics.** (**a**) *Right*, Voltage responses of a DG granule cell to 700 ms current pulses of amplitude varying from –25 pA to +25 pA (in steps of 5 pA), with normal ACSF (*black*) and in the presence of Riluzole in the bath (*blue*). *Left*, Input resistance ($R_{in}$) was calculated as the slope of the *V–I* plot depicting steady-state voltage response as a function of the injected current amplitude. (**b**) Voltage response of the example neuron to 5 alpha-current injections arriving at 20 Hz, depicting temporal summation. Temporal summation ratio ($S_\alpha$) was computed as the ratio of the amplitude of the fifth response to that of the first. (**c**) *Top,* Voltage response of the example neuron to a 700-ms current pulse of 250 pA in ACSF (*black*) and in the presence of riluzole (*blue*). *Bottom*, Frequency of firing plotted as a function of injected current amplitude for the example cell. Note that these are firing frequencies converted from the number of spikes obtained for 700-ms duration pulses. (**d**) *Left*, Voltage responses of the example neuron to the chirp current (**Fig. 1b**, top) in ACSF (*black*) and in the presence of Riluzole (*blue*). *Right*, Impedance amplitude computed from the current stimulus shown in **Fig. 1b** (*top*) and the voltage responses shown on the left. $|Z|_{max}$ represents the maximum impedance amplitude, $Q$ is resonance strength and resonance frequency is represented by $f_R$. (**e–l**) Population data of measurements from all DG granule cells recorded before and after adding riluzole to the bath: RMP (**e**); input resistance, $R_{in}$ (**f**); membrane potential dependence of input resistance (*G*); temporal summation (**h**); impedance amplitude, $|Z|_{max}$ (**i**) and its voltage dependence (**j**); firing rate at 0–250 pA current injection (**k**) and for 250 pA current injection (**l**). For panel **k**, statistical comparisons were performed with paired Student's $t$ test; *: $p < 0.05$.

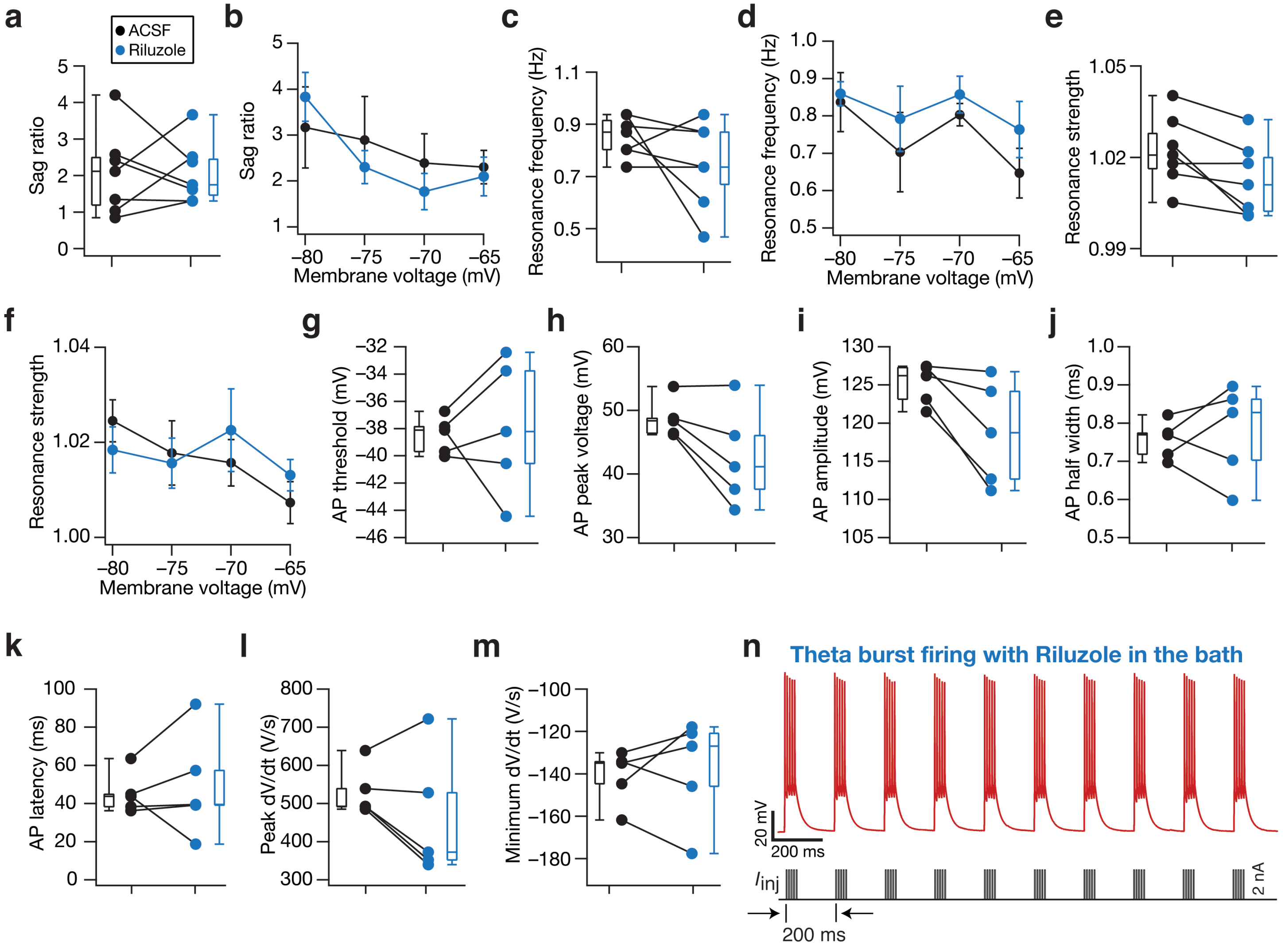


**Supplementary Figure S10. Impact of acute treatment with Riluzole, a persistent sodium channel blocker, on several sub-threshold and action potential measurements of DG granule cells.** (**a–m**) Population data of measurements from all DG granule cells recorded before (black) and after (blue) adding riluzole to the bath: Sag ratio (**a**) and its voltage dependence (**b**); resonance frequency (**c**) and its voltage dependence (**d**); resonance strength (**e**) and its voltage dependence (**f**); AP threshold (**g**); AP peak voltage (**h**); AP amplitude (**i**); AP half width (**j**); latency to first spike (**k**); peak (**l**) and minimum (**m**) dV/dt of the action potential waveform. Although there was a trend of consistent reduction in AP amplitude and AP peak voltage, none of the measurements depicted in this figure were statistically significant with Wilcoxon signed rank test. (**n**) Membrane voltage (*top*) recorded from the example granule cell in response to the theta-patterned current injection (*bottom*) in the presence of riluzole in the ACSF, showing 50 action potential elicited during a theta burst firing (TBF) pattern. Similar to other cases, this train was repeated thrice (total 150 action potentials) with 10 s inter-train intervals to form the TBF protocol.

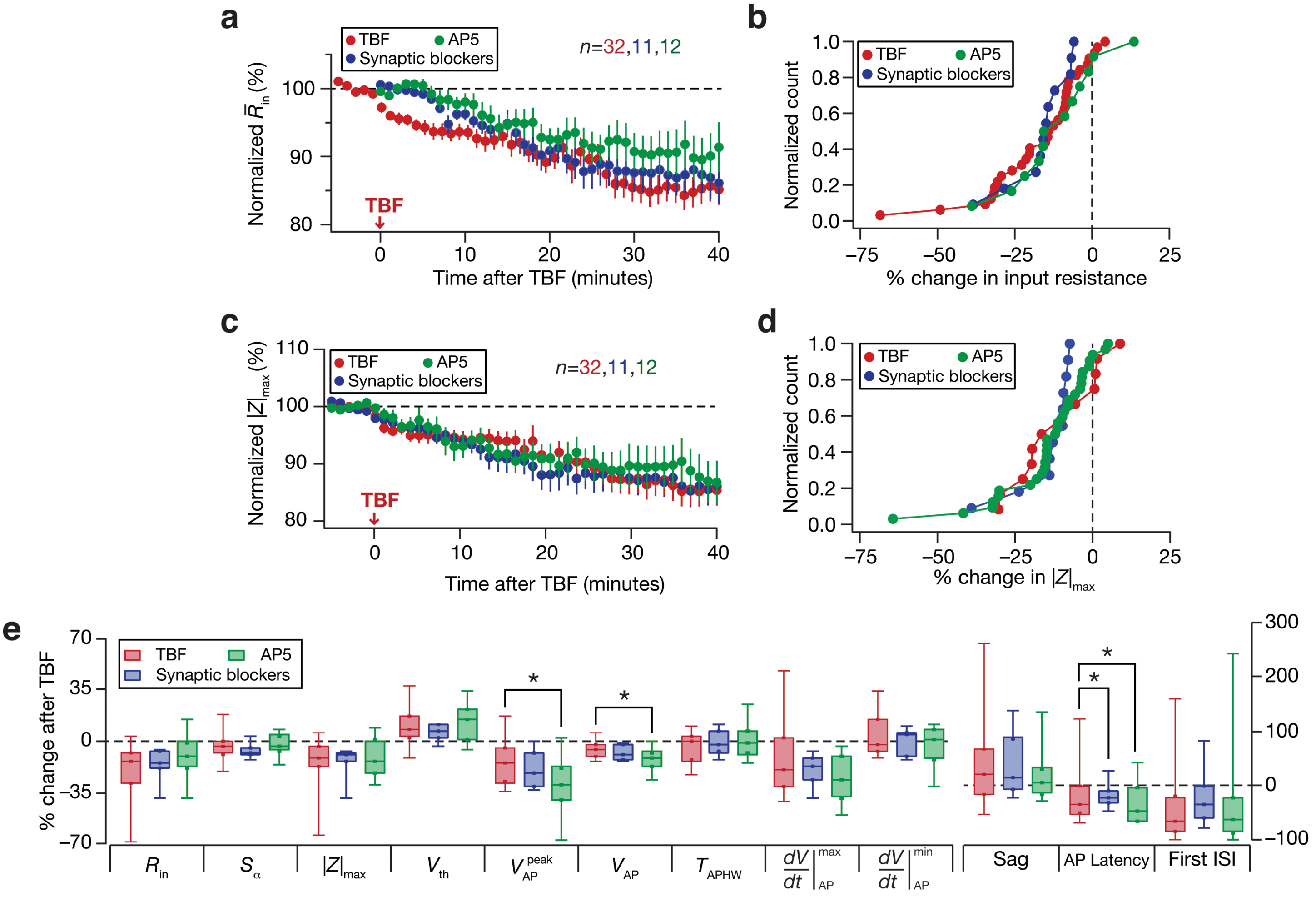


**Supplementary Figure S11. Activity-dependent intrinsic plasticity in DG granule cells was independent of synaptic receptors.** For all panels, the "Syn Block" (blue) group ($n$=11) corresponds to experiments where neurons were subjected the TBF protocol in the presence of synaptic blockers (AMPAR, $GABA_AR$ and $GABA_BR$). The "AP5" (green) group ($n$=12) corresponds to experiments where neurons were subjected the TBF protocol in the presence of AP5, an NMDAR antagonist. These are compared with the TBF group where normal ACSF was employed. (**a**) Temporal evolution of percentage change in input resistance estimate ($\bar{R}_{in}$; Mean ± SEM). (**b**) Normalized count of neurons from the three groups plotted as functions of percentage change in $R_{in}$ (data from **Fig. 9a**) (**c–d**) Same as (**a–b**), representing impedance amplitude ($|Z|_{max}$; **Fig. 9b**). (**e**) Plots comparing percentage change in various measurements from their initial values to end of experiment values. The list of symbols and corresponding measurements are enumerated in Supplementary Table S1. The Wilcoxon rank sum test was employed for $p$ value calculation to compare percentage changes in the control group *vs*. those in the TBF group.

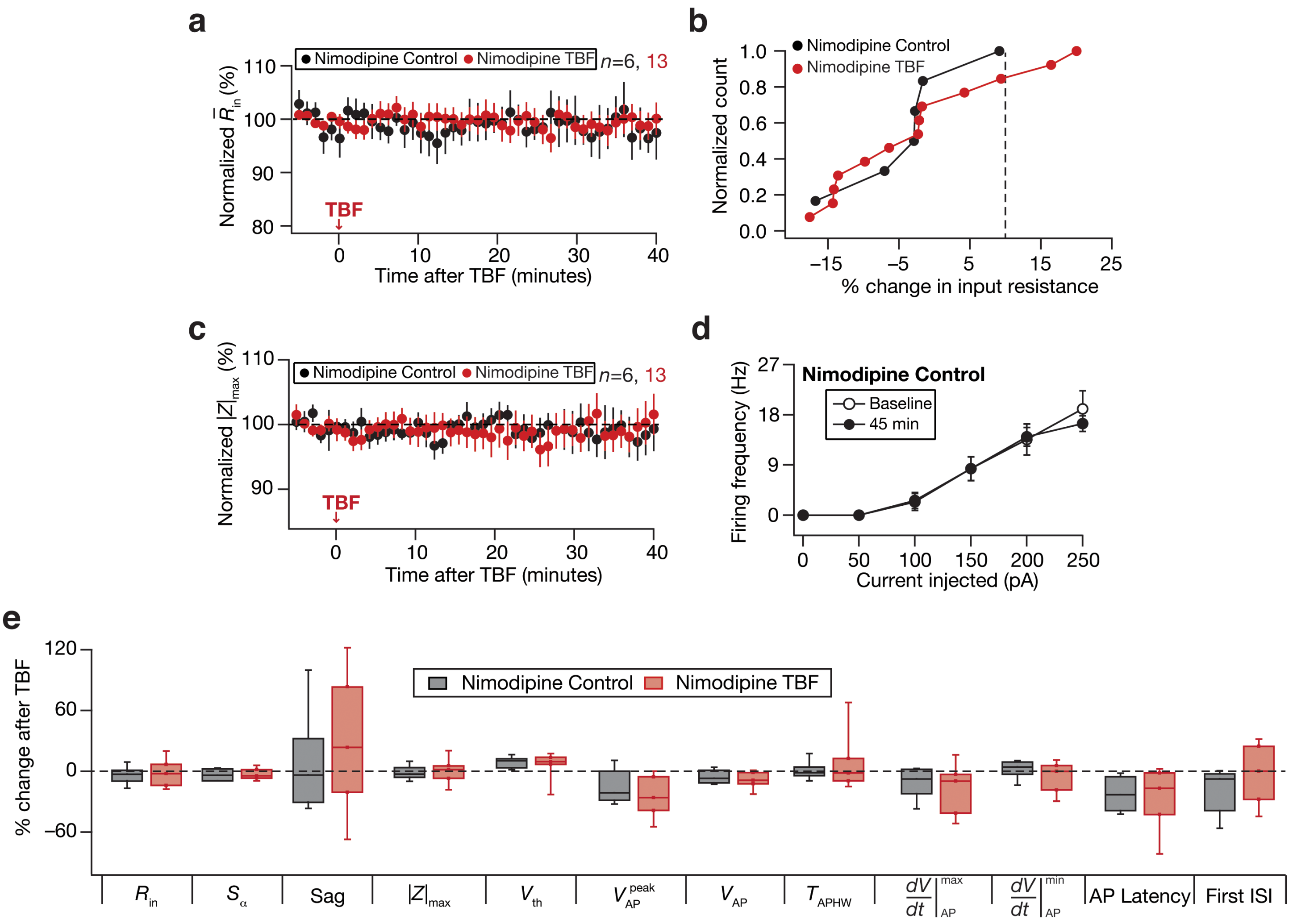


**Supplementary Figure S12. Activity-dependent intrinsic plasticity was blocked by Nimodipine, a *L*-type calcium channel blocker.** For all panels, the Nimodipine Control (black) group ($n$=6) corresponds to experiments where no protocol was applied through the 45-min period of the experiment, and Nimodipine TBF (red) group ($n$=13) is for neurons subjected to theta burst firing protocol. All experiments reported in this figure were performed in the presence of 10 μM Nimodipine in the ACSF bath solution. (**a**) Temporal evolution of percentage change in input resistance estimate ($\bar{R}_{in}$; Mean ± SEM). (**b**) Normalized count of neurons from **Fig. 9d** plotted as a function of percentage change in $R_{in}$. (**c**) Temporal evolution of percentage change in maximal impedance amplitude ($|Z|_{max}$). (**d**) Frequency of firing plotted as a function of injected current amplitude for the Nimodipine control group, where no TBF protocol was applied. (**e**) Plots comparing percentage change in various measurements from their initial values to end of experiment values. The list of symbols and corresponding measurements is enumerated in Supplementary Table S1. The Wilcoxon rank sum test was employed for $p$ value calculation to compare percentage changes in the control group *vs*. those in the TBF group.

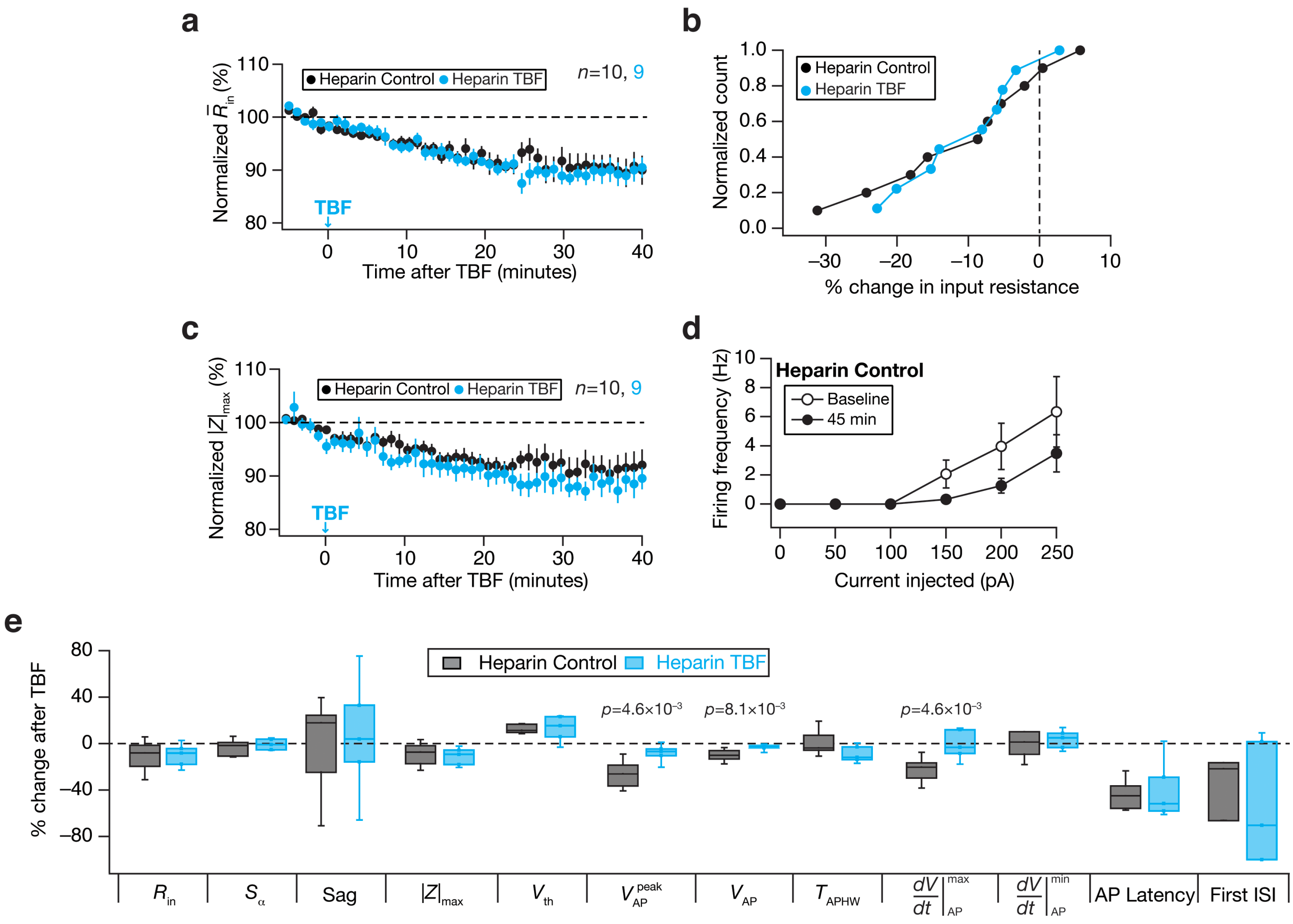


**Supplementary Figure S13**. **Activity-dependent intrinsic plasticity was blocked by Heparin, an InsP$_3$R blocker.** For all panels, the Heparin Control (black) group ($n$=6) corresponds to experiments where no protocol was applied through the 45-min period of the experiment, and Nimodipine TBF (cyan) group ($n$=13) is for neurons subjected to theta burst firing protocol. All experiments reported in this figure were performed in the presence of 1 mg/mL Heparin in the pipette. (**a**) Temporal evolution of percentage change in input resistance estimate ($\bar{R}_{in}$; Mean ± SEM). (**b**) Normalized count of neurons from **Fig. 9g** plotted as a function of percentage change in $R_{in}$. (**c**) Temporal evolution of percentage change in maximal impedance amplitude ($|Z|_{max}$). (**d**) Frequency of firing plotted as a function of injected current amplitude for the Heparin control group, where no TBF protocol was applied. (**e**) Plots comparing percentage change in various measurements from their initial values to end of experiment values. The list of symbols and corresponding measurements is enumerated in Supplementary Table S1. The Wilcoxon rank sum test was employed for $p$ value calculation to compare percentage changes in the control group *vs*. those in the TBF group.

**Supplementary Table S1: Sub- and supra-threshold physiological properties measured before and 40 minutes after TBF.** These statistics correspond to the data plotted in Fig. 2. $p$ values correspond to paired Student's $t$ test, and data are represented as mean ± SEM.

| | Measurement (units) | Before TBF | After TBF | Significance |
|---|---|---|---|---|
| **Sub-threshold measurements** | | | | |
| 1 | Resting membrane potential, $V_{RMP}$ (mV) | –75.9 ± 0.6 | –72.3 ± 1.2 | $p$=6.3×10$^{-4}$ |
| 2 | Input resistance, $R_{in}$ (MΩ) | 141.8 ± 7.5 | 118.2 ± 8.02 | $p$=4.4×10$^{-6}$ |
| 3 | Maximal impedance amplitude, $\|Z\|_{max}$ (MΩ) | 155.9 ± 8.2 | 134.2 ± 8.8 | $p$=1.4×10$^{-5}$ |
| 4 | Resonance frequency, $f_R$ (Hz) | 0.77 ± 0.01 | 0.99 ± 0.20 | $p$ = 0.27 |
| 5 | Resonance strength, $Q$ | 1.03 ± 0.03 | 1.03 ± 0.003 | $p$ = 0.86 |
| 6 | Total inductive phase, $\Phi_L$ (rad.Hz) | 0.01 ± 0.002 | 0.01 ± 0.001 | $p$ = 0.33 |
| 7 | Sag (%) | 2.6 ± 0.14 | 3.3 ± 0.31 | $p$=0.037 |
| 8 | Summation ratio of αEPSPs, $S_\alpha$ | 1.3 ± 0.03 | 1.23 ± 0.02 | $p$=3.5×10$^{-3}$ |
| **Supra-threshold measurements** | | | | |
| 1 | Firing frequency | Significant increase after TBF | | Fig. 2$F$ |
| 2 | AP threshold, $V_{th}$ (mV) | –38.9 ± 0.6 | –42.9 ± 1.4 | $p$=5.4×10$^{-4}$ |
| 3 | AP peak, $V_{AP}^{peak}$ (mV) | 51.0 ± 1.7 | 43.2 ± 1.7 | $p$=2.2×10$^{-5}$ |
| 4 | AP amplitude, $V_{AP}$ (mV) | 126.2 ± 1.8 | 118.4 ± 1.9 | $p$=2.0×10$^{-5}$ |
| 5 | AP half-width, $T_{APHW}$ (ms) | 0.85 ± 0.02 | 0.82 ± 0.02 | $p$=0.17 |
| 6 | Peak dV/dt, $\left.\frac{dV}{dt}\right\|_{AP}^{max}$ (V/s) | 528.5 ± 22.7 | 443.5 ± 20.8 | $p$=9.6×10$^{-4}$ |
| 7 | Min dV/dt, $\left.\frac{dV}{dt}\right\|_{AP}^{min}$ (V/s) | –125.2 ± 3.3 | –128.3 ± 3.6 | $p$=0.31 |
| 8 | Latency to first spike, $T_{1AP}$ (ms) | 54.7 ± 5.6 | 49.2 ± 10.7 | $p$=0.44 |
| 9 | First ISI, $T_{1ISI}$ (ms) | 38.1 ± 9 | 8.5 ± 1.62 | $p$=5.0×10$^{-3}$ |
